# Molecular Insights into the Collagen-binding Specificity of Discoidin Domain Receptors

**DOI:** 10.64898/2026.10.13.758775

**Authors:** Miriam Prinzler, Lukas Jaegers, Jean-Daniel Malcor, Birgit Leitinger, Abhishek A. Jalan

**Author notes:** Equal contribution (Miriam Prinzler, Lukas Jaegers).

## Abstract

Discoidin domain receptors, DDR1 and DDR2, are a subfamily of homologous receptor tyrosine kinases that are activated by the extracellular matrix (ECM) collagens. Together the two DDRs recognize at least seven of the twenty-eight known collagen types (Col1-28) with distinct specificities. For example, Col1-3 and Col5 that exist as fibres in connective tissues activate both DDRs whereas Col6 microfilaments found in ECM of skeletal muscles preferentially activates DDR1. Further, Col4 found in basement membrane as mesh-like networks preferentially activates DDR1 but similarly network-forming Col10 abundant in cartilage preferentially binds and activates DDR2. These preferential interactions activate specific signalling pathways driving distinct physiological processes and pathologies, yet their molecular basis is currently not understood. Here, we show that the collagen-binding specificity of the DDRs is determined by electrostatic interactions at the receptor-ligand binding interface. The collagen-binding site in the two DDRs differ in the nature and placement of electrostatically charged residues. The known DDR-binding epitopes in Col1-Col3 also contain charged residues positioned to interact with those on the DDRs. We hypothesize that these interfacial electrostatic interactions between the DDRs and collagens guide their recognition specificity. We experimentally test this hypothesis and show that triple-helical peptides (THPs) containing rationally placed charged residues have biased affinities for the two DDRs. Aside from advancing mechanistic understanding of the different collagen-DDR interactions, our results can facilitate better understanding of the associated signalling pathways and the design of therapeutically active DDR-specific antagonists.

## Introduction

Cells sense their microenvironment via surface proteins that transduce biochemical cues present in the extracellular matrix (ECM).^1^ Cellular response to these cues regulate migration, proliferation and differentiation while dysregulated signalling contributes to numerous diseases. As the most abundant component of the ECM, collagens present several biochemical and mechanical cues.^2,3^ They interact with several families of mammalian cell surface receptors, including integrins^4^, discoidin domain receptors (DDR1 and DDR2)^5,6^, glycoprotein VI^7,8^, leukocyte-associated immunoglobulin-like receptor^9,10^ and proteins of the mannose receptor family^11^ to regulate a variety of cellular functions. Among the receptors, DDR1 and DDR2 have recently attracted significant attention due to their role in fibrosis^12^, cardiovascular diseases^13^, osteoarthritis^14,15^ and several forms of cancer^16^.

DDR1 and DDR2 constitute a subfamily of homologous proinflammatory receptor tyrosine kinases (RTK) broadly comprising an intracellular kinase, a transmembrane and an extracellular region^17^. Collagens bind to the extracellular region, which results in receptor activation, autophosphorylation of the intracellular kinase domain and downstream signalling events.^18,19^ On a molecular level, collagens are characterized by a rod-like structure called a triple-helix composed of three polypeptides.^20^ The polypeptides themselves contain a repetitive sequence (G-X-Y)_n_, where every third amino acid is glycine and *X-* and *Y-*positions are frequently non-glycine residues. Human collagen genes encode forty-four polypeptides referred to as *α*-chains that assemble into twenty-eight distinct types of collagens (Col1-28), which exist as fibres, networks or filaments in the ECM.^2^ Binding and activation experiments conducted with tissue-derived collagens suggest that several different collagen types recognize the two DDRs but some preferentially recognize either DDR1 or DDR2.^17,21,22^ For example, fibrillar collagens Col1-3 and Col5 found in connective tissues activate both DDRs but the network-forming Col4 present in the basement membrane preferentially activates DDR1^21,23^. Conversely, Col10, also a network-forming collagen that facilitates cartilage formation and its subsequent ossification into bone tissues^24^, preferentially binds and activates DDR2.^25^ Further, the microfilament-forming Col6, which is found in the ECM of skeletal muscles and plays a prominent role in chondrocyte proliferation^26^, protection of neurons against Amyloid β toxicity^27^ and Schwann cell differentiation^28^, preferentially activates DDR1.

The molecular basis of the collagen-binding preference of DDRs is currently unknown, yet it profoundly affects the associated physiological functions. For example, DDR1-Col1 interaction is implicated in the organization and remodelling of fibrillar collagen matrices surrounding the cells.^29,30^ Aberrant DDR1-Col1 interaction can cause pathological deposition of collagen, promoting fibrosis, or insufficient matrix organization, causing pathological cell proliferation, which is a hallmark of invasive cancer. Thus, DDR1-Col1 interaction represents an antifibrotic as well as anti-cancer target.^31^ In a different context, mutations in Col4 drive Alport Syndrome, a debilitating kidney disease.^32^ Notably, genetic loss of DDR1 in mice deficient in a Col4 *α*-chain partially restores renal function,^33^ suggesting that DDR1-Col4 signalling contributes directly to disease pathogenesis. Therefore, inhibition of DDR1-Col4 interaction may offer a therapeutic strategy for the treatment of Alport Syndrome. DDR1 also plays a prominent role in atherosclerosis^34,35^ where it promotes smooth muscle cell migration and proliferation^36^, resulting in progressive fibrosis^37^ and calcification^38^ of the atherosclerotic plaque. Vascular cells express several DDR1-binding collagens, including Col1 and Col3-Col6.^39^ Thus, specific inhibition of their interaction with DDR1 could slow progression of atherosclerosis.

DDR2 provides an equally compelling case for dissecting the collagen-binding specificity of DDRs. DDR2 activation induces matrix metalloproteinase (MMP) −1 expression, which proteolytically remodels the ECM and regulates matrix turnover.^14^ Thus, preferential interaction of DDR2 with fibrillar collagens Col2 and Col3 plays a key role in regulating ECM remodelling by fibroblasts and chondrocytes.^40^ Dysfunction within this DDR2-mediated signalling pathway contributes to fibrosis of organs and promotes tumour invasion by facilitating collagen degradation and stromal remodelling.^41^ Additionally, DDR2 upregulation in chondrocytes results in MMP-13 mediated degradation of collagen^42–44^ resulting in osteoarthritis while reduction of DDR2 expression slows cartilage degradation.^45^ Further, inhibition of DDR2 binding to Col2 (the most abundant collagen type in cartilage) and Col10 (a marker for hypertrophic chondrocytes that is upregulated in osteoarthritis)^38^ slows progression of osteoarthritis^15^. Finally, loss-of-function mutations in DDR2 causes dysfunctional interaction with collagen, which impairs chondrocyte proliferation and bone growth during spondylo-meta-epiphyseal dysplasia (SMED).^46^

Together, these examples underscore that the specificity with which DDRs recognise the different collagens is central to their function and that understanding its molecular basis is the first key step for therapeutic targeting of these interactions. Here, we present theoretical analyses and experimental data that suggest that the collagen-binding specificity of DDRs is determined by electrostatic interactions. Specifically, we show that the interaction between electrostatically charged residues at the collagen-binding interface of the DS-domains as well as the bound collagen ligand contribute to their binding specificity. We also designed triple-helical peptides (THPs) containing electrostatically charged residues that show biased affinities for DDR1 and DDR2. In the process, we obtained a THP that preferentially recognizes DDR2 over DDR1. Aside from advancing mechanistic understanding of the collagen-binding specificity of DDRs, our results could facilitate future studies into DDR-specific signalling pathways, how these drive pathology and strategies for receptor-specific therapeutic interventions.

## Results

### The DS-domains of DDR1 and DDR2 differ in electrostatic charges

The extracellular region of DDRs is composed of two tandem domains that bear structural homology to the discoidin (DS) domain of the amoeba *Dictyostelium discoideum*.^47^ The DS-domain distal to the cell membrane recognizes collagens via a consensus binding epitope RGQOGVMGFO (O = 4(R)-hydroxyproline, hereafter also abbreviated as Hyp) conserved in Col2 and Col2.^48–50^ Our analysis of the isoelectric point (pI) and the surface electrostatic potential of the collagen-binding DS-domains as well as the distribution of electrostatically charged residues at the ligand binding interface reveals marked differences between DDR1 and DDR2:

a. pI of the human DDR1 DS-domain is 7.7 while that of DDR2 is 5.3 (**see methods**). Thus, at physiological pH, the DS-domain of DDR1 is largely neutral (*z* = +0.5) while that of DDR2 carries a net negative charge of −7.3 (**supplementary fig. 1**). The shift in charge of ∼8 units suggests different global electrostatic character of the two DS-domains.
b. Analysis of the surface electrostatic potential of the DS-domains using Adaptive Poisson-Boltzmann Solver (APBS)^51^ suggests differences near the collagen binding site of the DS-domains **(Figure 1A, B)**. The DDR2 DS-domain displays a negative surface potential in and around the deep binding pocket that accommodates the Phe of the ligand RGQOGVMGFO epitope. This binding pocket has comparatively less negative potential in DDR1. Further, the Arg of the epitope points towards a negatively charged patch of the DDR2 DS-domain that has previously been proposed to indicate a long-range electrostatic interaction^52^. The surface potential of this patch in DDR1 is comparatively less negative. Additionally, the C-terminal end of the bound ligand is positioned close to a largely neutral surface potential in the DDR2 DS-domain but, in the case of DDR1, this surface bears a strongly positive charge.
c. The co-crystal structure of the DDR2 DS-domain bound to a THP containing the RGQOGV(Nle)GFO epitope (Nle = norleucine, a bioisostere of Met introduced to increase binding affinity) reveals that Phe binds an amphipathic pocket lined by five DDR2 residues that are strictly conserved in DDR1 (**Figure 1B, C**).^52^ However, five other residues – H110, I112, H172, S173 and N175 – that peripherally contribute to the binding interface vary between the two DDRs.^53^ Leitinger and co-workers hypothesized that these peripheral residues might contribute to the observed collagen-binding specificity of the two DDRs. To probe this, they mutated the five DDR2 residues to those in DDR1 (H110L, I112K, H172R, S173V and N175S). The resulting hybrid DDR2 construct named DDR2-H3, contrary to its native specificity, recognised Col4.^53^ Of the five mutations, I112K and H172R alter the electrostatic charge profile close to the collagen binding site.
d. The DS-domain of DDR2 contains two anionic patches located at diametrically opposite ends of the ligand-binding site **(Figure 1C**). The first anionic patch created by residues E66, E67 and D69 of DDR2 loses its negative charged density in DDR1 due to the substitution of E66 with S67. The second patch formed by E79, D81 and D82 residues in DDR2 is also less negative in DDR1 due to the substitution of E79 and D81 with F80 and K82, respectively. Importantly, the DDR1 DS-domain contains a constellation of positively charged residues, K112, R116 and R172, lining one side of the collagen-binding site. In the case of DDR2, the equivalent sequence positions are occupied by electrostatically neutral H110, I112 and P116.

**Fig. 1.**
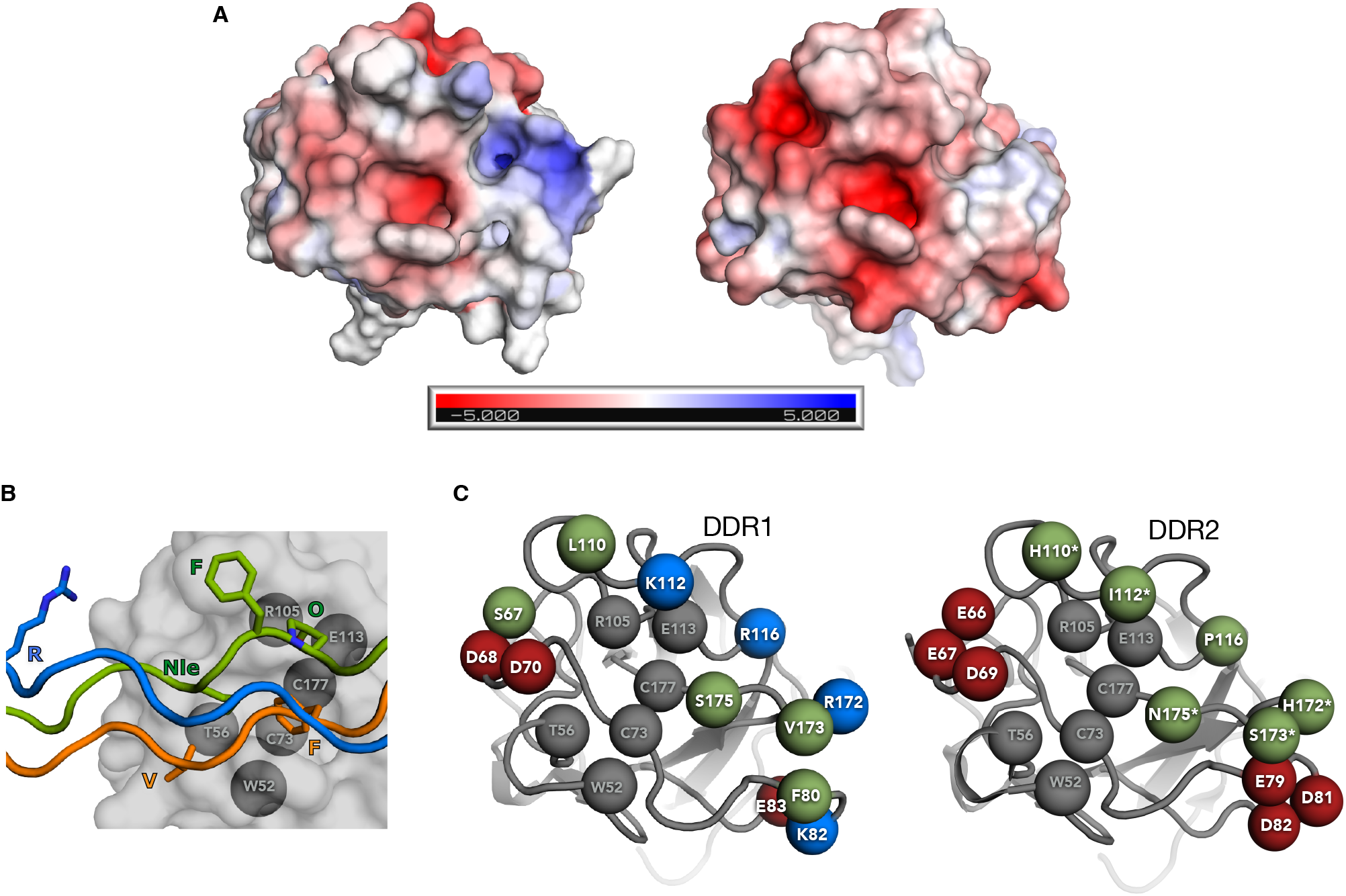
Electrostatic charge at the collagen-binding face of DS-domains. **(A)** Comparison of the surface electrostatic potential at the THP-binding face of the DS-domains of DDR1 (left, pdb:4ag4)^54^ and DDR2 (right, pdb:2wuh^55^; the bound THP is not shown for clarity). Electrostatic surface potentials were calculated using APBS^51^ and are coloured from −5 *k*_b_Tec^-1^ (red) to +5 *k*_b_Tec^-1^ (blue), where *k*_b_ is Boltzmann constant, T is temperature and e is electronic charge **(B)** Previously published co-crystal structure of THP bound to the DS-domain of DDR2 (pdb:2wuh)^52^ highlighting interfacial interactions. Polypeptides in a triple helix are staggered by one amino acid creating a leading (green), middle (orange) and trailing (blue) chain position. Six THP residues of the DDR-recognizing motif GARGQOGV(Nle)GFO [O= 4(R)-hydroxyproline; Nle = norleucine, a bioisostere of Met] spread across the three chains contribute to the interfacial interaction: leading chain Phe and Nle, middle chain Phe and Val and trailing chain Arg. C*α*-atoms of residues critical for interaction are shown as grey spheres. (**C**) Positive (blue), negative (red) and neutral (green) residues located near the binding site and differing between the two DS-domains are also highlighted. The five residues in the DDR2 DS-domain previously mutated to reverse its collagen-binding specificity^49^ are marked with asterisk. C*α*-atoms of residues in DDR1 and DDR2 DS-domains critical for collagen recognition and conserved across both DDRs are shown as grey spheres.

Collectively, the above observations suggest that the DS-domains of DDR1 and DDR2 differ substantially with respect to the distribution of electrostatic charges globally but, most prominently, near the collagen-binding site.

### Electrostatically charged residues in the collagen ligand determine DDR-recognition specificity

In the previous section, we focused on differences in the electrostatic charge profile of the DDR1 and DDR2 DS-domains. In this section, we analyse previously published data to show that charged residues on the THP ligand guide preference for DDR1 or DDR2. Native collagens are highly aggregation-prone under laboratory conditions, which complicates direct molecular study of their interaction with DDRs. Farndale and co-workers circumvented this challenge by designing a library of THPs where short overlapping fragments of native collagen sequences are included as ‘guests’ into a (GPP)_5_ ‘host’. The guest sequences contain potential epitopes for recognition of other proteins whereas the host sequence promotes triple-helical structure.^56^ Scanning each THP in this ‘toolkit’ library for binding activity reveals the epitope capable of recognizing a given protein.^57^ The currently available Col2 and Col3 toolkits contain 56 and 57 THPs, respectively, spanning their entire collagen domains. DDR1 and DDR2 recognize a highly restricted subset of THPs in both toolkits.^49,50^ Analysis of these THPs suggest that the placement and type of electrostatic charge on the bound ligand contribute to their DDR-binding specificity.

As shown in **Table 1**, DDR2 binds multiple toolkit THPs containing the GxyGFO motif where GAO, GAT, GER, GIO and GAA are tolerated in place of Gxy.^49,50^ Further, THPs containing Leu (II-5; II-46) and Met (III-44) in place of Phe also bind DDR2, suggesting a fairly promiscuous ligand binding site. However, DDR1 is more restrictive, as reflected in its inability to bind THPs where the minimal recognition motif deviates from GVMGFO (II-5, II-13, II-39, II-44, III-5, III-39, III-44).^49^ A notable exception is THP II-46 that contains GPOGLT and binds DDR1 but not DDR2. Aside from the minimal GVMGFO motif, sequence variations in regions flanking it also strongly influence specificity. For example, the GVMGFO motif conserved in Col2 and Col3 is flanked by an Arg two triplets towards the N-terminal and a Lys in the triplet immediately towards the C-terminal. THP II-22 lacking the C-terminal Lys recognizes both DDRs but THP II-23 lacking the N-terminal Arg recognizes only DDR2.^49^ Thus, the N-terminal Arg is required for DDR1 recognition but not DDR2 and the C-terminal Lys is dispensable for recognition of both DDRs. As noted before, in the crystal structure of DDR2 DS-domain bound to THP^52^, Arg points towards the acidic patch created by residues E66, E67 and D69 of the DS-domain. This long-range electrostatic interaction is likely weekend in DDR1 due to substitution of E66 with S67. Consequently, DDR1 is more sensitive to the loss of Arg-mediated electrostatic interaction. It is noteworthy that the variant construct DDR2-H3, which contains the H110L mutation close to this acidic patch does not bind THPs II-23 and III-39 that lack the N-terminal Arg, thus mimicking the behaviour of DDR1 (**Table 1**).^49^ In the DDR2-THP co-crystal structure, H110 interacts with Phe of the ligand via an apparent π-π stacking interaction between their aromatic rings^52^. The H110L mutation abrogates this interaction. Thus, the N-term Arg is likely required as compensating interaction for binding DDR1 but not DDR2. This further emphasise the sophisticated role played by the N-terminal Arg in determining the specificity of the collagen ligand for DDR1 and DDR2.

**Table 1.**
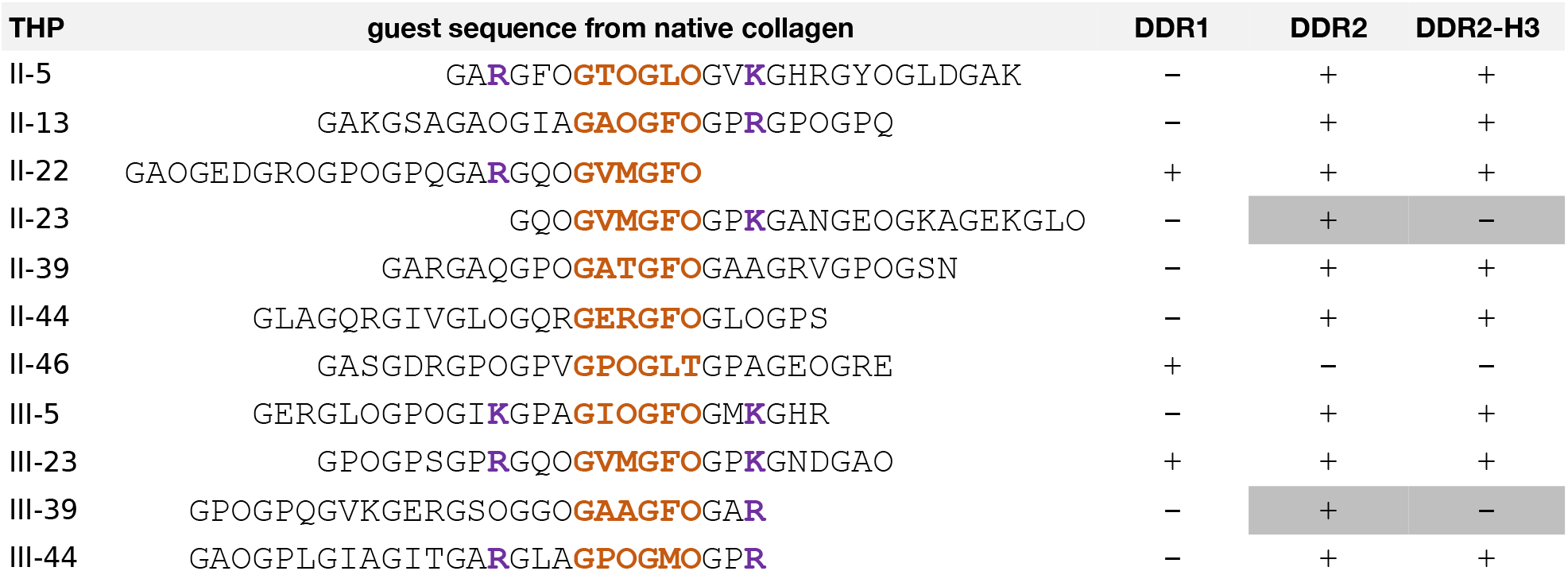
Binding of toolkit THPs containing guest sequences from Col2 or Col3 to DDR1, DDR2 or its mutant construct DDR2-H3 containing H110L, I112K, H172R, S173V and N175S mutations, as published previously^49,50^. Hydroxyproline residues are indicated as “O”. Positive and negative binding interactions are denoted as “+” and “-”, respectively. The minimal DDR-binding motif in the THP guest sequence is shown in orange and the electrostatically charged residues that appear to contribute to the specificity of binding to DDR1 or DDR2 are shown in purple. THPs with opposite preferences for DDR2 and DDR2-H3 are shaded grey.

### Insights into the DDR-recognition preference from Col4 and Col10

Col4 preferentially recognizes DDR1^5,6^ whereas Col10 preferentially recognizes DDR2^58^. However, the corresponding DDR-binding sites remain unknown. We identified potential DDR1-binding sites in Col4 and DDR2-binding sites in Col10 and examined these for features that might account for their opposite DDR recognition preferences. We did not pursue experimental validation of these sites in this study due to design challenges – Col10 is a homotrimeric collagen^59^ i.e. its triple-helical domain is composed of identical polypeptides. However, Col4 is a subfamily of collagens containing three distinct heterotrimers that vary in the composition of the triple helix^60^. Design of homotrimeric THPs is straightforward as peptides containing sufficient G-X-Y repeats self-assemble into a triple helix^61^. In contrast, the heterotrimer design is complicated by the combinatorically distinct triple helices that could assemble in a mixture of two or more unique peptides.^62,63^ Therefore, due to significant challenges associated with the design of heterotrimers, a comparative assessment of the binding of Col4 and Col10 sites to DDR1 and DDR2 is non-trivial and will be pursued in future.

The previously published crystal structure of the THP containing GA**R**GQO**GV(Nle)GFO** epitope bound to the DS-domain of DDR2 identified six collagen residues that primarily determine the binding interaction: Phe, Nle and Hyp of the leading chain, Phe and Val of the middle chain and Arg of the trailing chain (**Figure 1B**).^52^ Subsequently, alanine mutagenesis with a THP containing GA**R**GQO**GVMGFO** revealed that Met and Phe are indispensable for recognition of both DDRs whereas mutation of Val or Hyp decreases affinity.^49,50^ There is conflicting data on how Arg influences recognition. As shown in **Table 1**, THPs lacking the N-terminal Arg recognize DDR2 but not DDR1. However, mutagenesis experiments find that mutation of Arg to Ala only marginally decreases affinity for the two DDRs.^49,50^ Crucially, THPs lacking the N-term Arg, and thus unable to recognize DDR1, contain GAOGFO, GATGFO etc. in place of the GVMGFO motif. As noted before, it is likely that these biochemically similar motifs facilitate weaker binding than the GVMGFO and thus require Arg as a compensating interaction. Since Col4 or Col10 α-chains do not contain the GVMGFO motif, we here presume that Arg is required for recognition of DDR1 but not DDR2. Further, in the crystal structure, the leading chain Phe interacts with a H110 of the DDR2 DS-domain via π-π stacking of their aromatic rings and the middle chain Phe sits in the amphipathic binding pocket.^52^ It is currently not clear if both Phe’s are critical for binding. We have previously reported that mutation of the trailing chain Phe of the DDR-binding epitope in Col1 is not required for recognition.^64^ Thus, we also presume here that both the leading and middle chain Phe are required for recognition while the trailing chain Phe is dispensable. To summarize, these observations suggest that the Phe of the leading and middle chain, Met of the leading chain and Arg of the trailing chain are minimally required for recognition of DDR1 while Arg is dispensable for DDR2 recognition (**Fig. 2A**). These residues constitute a sequence-based constraint for DDR recognition. Since a collagen triple helix contains three staggered ɑ-chains related by approximately 120° rotation along the helical axis, the residues constituting the sequence constraint can be presented in three symmetry-equivalent registers (**Fig. 2B**). In the crystal structure, Phe of the middle chain sits in a pocket on the DS-domain^52^. Thus, the alternative registers are generated by rotating the triple helix by ∼120° along the helical axis, such that the equivalent Phe residue from the leading or trailing chain occupies the same pocket while the remaining critical residues occupy the corresponding symmetry-related positions.

**Fig. 2.**
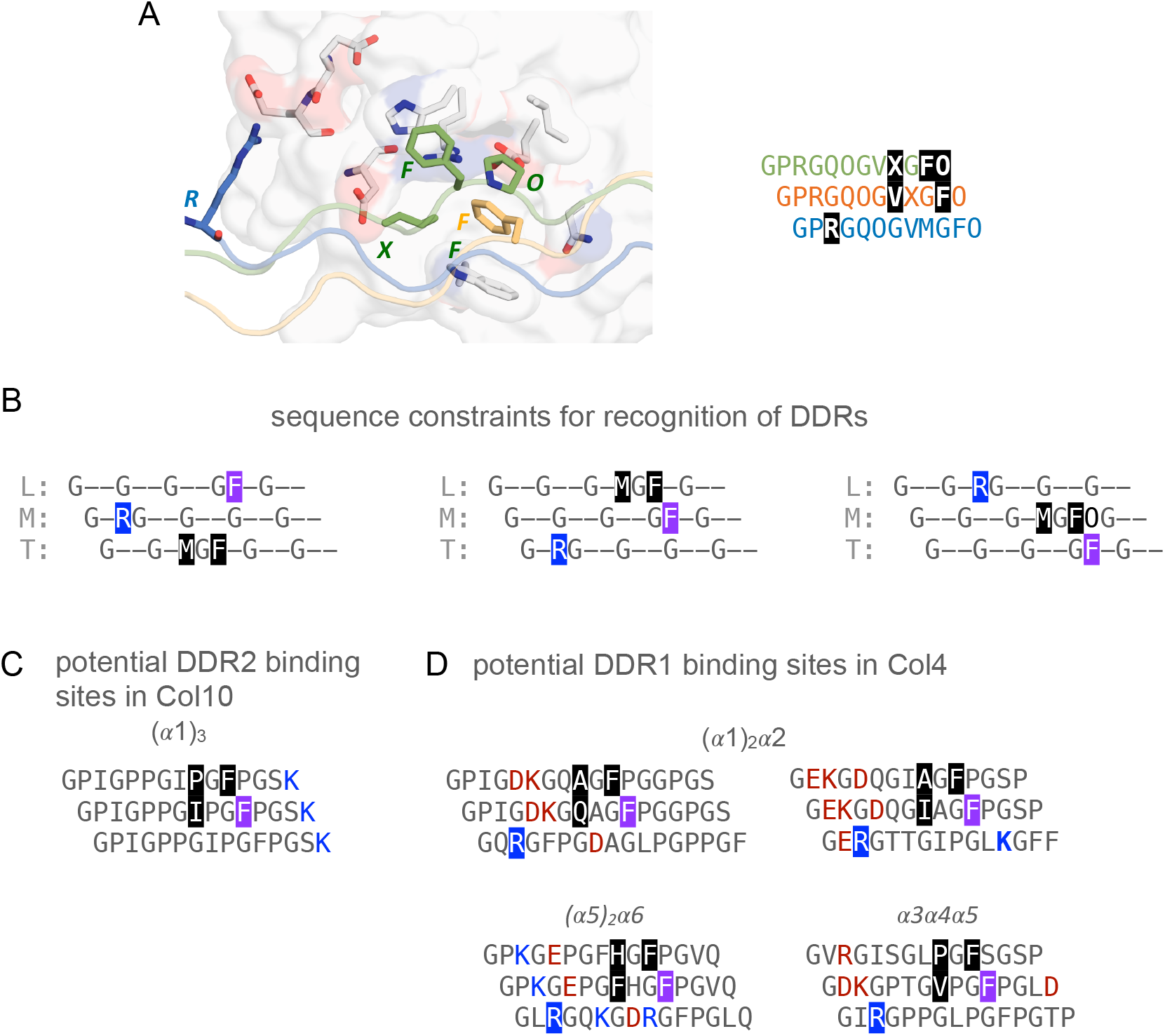
Potential DDR binding sites in Col10 and Col4. **(A)** Previously published co-crystal structure^52^ highlighting the THP residues (shaded grey) from the leading (green), middle (orange) and trailing (blue) chains that primarily contribute to binding interactions with the DDR2 DS-domain**. (B)** A minimal set of sequence constraints identified for the recognition of DDRs in the three orientations of the triple helix**. (C)** Potential DDR2 recognition site identified in Col10 homotrimer. **(D)** Potential DDR1 recognition sites in Col4 heterotrimers. Phe that could potentially bind the amphipathic pocket on the DS-domains is shaded purple, Arg interacting with the electrostatic patch is shaded blue and auxiliary interfacial interactions are shaded black. Positively and negatively charged residues surrounding the minimal motif are coloured red and blue, respectively.

We obtained human collagen sequences from UniProtKb database [*α*1(X)-Q9Y6Z7; *α*1(IV) – P02462; *α*2(IV) – P08572; *α*3(IV) – Q01955; *α*4(IV) – P53420; *α*1(V) – P29400; *α*1(VI) – Q14031], aligned them in JalView^65^, as published previously for homotrimer (*α*1)_3_^66^ of Col10 and the (*α*1)_2_*α*2, (*α*5)_2_*α*6 and *α*3*α*4*α*5 heterotrimers of Col4^67^ (**supplementary fig. 2**). We then searched for sites that match any of the three sequence constraints using a custom Python script that strictly required Phe but accepted residues Ala, Leu, Ile, Val, Pro or His instead of Met. The script also applied a post-filter to remove motifs that do not conform to strict Gly-X-Y repeat pattern. Deviations from this perfect pattern are called interruptions^68^, which structurally perturb the triple-helical backbone and cause local unwinding^69^. We excluded sites containing interruptions as these are unlikely to conform to the triple-helical structure required for binding to the DS-domains. The sequence search resulted in a single site GPIGPP**GIPGFP**GS**<u>K</u>**GD (Col10 residues 452-468) for Col10 (**Fig. 2C**), which is similar to the Col3 motif GIKGPA**GIOGFO**GM**<u>K</u>**GH found in THP III-5 (**Table 1)** that only binds DDR2^49^. Importantly, this potential Col10 motif and THP III-5 share a Lys in the triplet immediately C-terminal to the consensus motif. As discussed in the later sections, Lys influences the DDR-binding specificity of THPs.

Previously published data suggests that all three Col4 heterotrimers -(*α*1)_2_*α*2, (*α*5)_2_*α*6 or *α*3*α*4*α*5-can recognize DDR1. For example, Vogel et al showed DDR1 binding activity for Matrigel Col4^5^ extracted from the basement membrane of Engelbreth-Holm-Swarm (EHS) sarcoma^70^, which exclusively contains the *α*1(IV) and *α*2(IV) chains. Further, two separate studies have shown that DDR1 broadly signals through the *α*5(IV) chain^71^ and specifically through the *α*3*α*4*α*5 heterotrimer^72^. This leaves ambiguity regarding the ability of the (*α*5)_2_*α*6 heterotrimer to act as a DDR1 ligand. Nonetheless, we searched all three heterotrimer alignments shown in **supplementary fig. 2** for potential DDR1 binding sites using our sequence constraints. This revealed two potential binding sites for the (*α*1)_2_*α*2 heterotrimer and one potential binding site each for the (*α*5)_2_*α*6 and *α*3*α*4*α*5 heterotrimers (**Fig. 2D**). In addition to the residues defined by the sequence constraint, all potential binding sites in Col4 contain charged residues surrounding the minimal DDR1-binding motif.

The binding mode predicted for these sites appears similar to that observed in the DDR2-THP crystal structure^52^ i.e. the middle chain Phe occupies the amphipathic pocket of the DS-domain. Thus, we used the crystal structure as a template to understand if the charged residues in the potential binding sites could interact with those on the DS-domains of DDR1 and DDR2 based on spatial proximity. As shown in **supplementary fig. 3**, the C-terminal Lys of the potential binding site in Col10 is proximal to the acidic patch created by E79, D81 and D82 of the DDR2 DS-domain. The distance between this Lys and the acidic patch precludes direct salt bridge contact, but a long-range electrostatic interaction is plausible. As noted before, this patch in DDR1 loses much of its negative charge due to substitutions with F80 and K82, which would weaken the electrostatic interaction with the Lys of the Col10 site. This could explain Col10 preference for DDR2 over DDR1. The interfacial electrostatic interactions between the Col4 (*α*1)_2_*α*2 and (*α*5)_2_*α*6 are presently difficult to pinpoint. The potential sites identified in both heterotrimers contain several positive and negatively charged residues in the vicinity of the N-terminal Arg of the minimal motif. These charges could influence the magnitude of long-range electrostatic interaction to the DDR1 and DDR2 DS-domains and bias affinity. However, this can only be revealed by experimental investigation of the interaction between DDRs and the respective heterotrimers. The potential site identified in *α*3*α*4*α*5 also contains several charged residues adjacent to the N-terminal Arg, which could modulate electrostatic interactions. Intriguingly, it also contains a C-terminal Asp that can form direct salt-bridge with R172 of the DDR1 DS-domain. Importantly, the DDR2 DS-domain cannot form this salt bridge due to presence of H172 at this sequence position.

### Rational design of THPs to understand how charged residues influence DDR recognition

We designed THPs containing the motif GP**R**GPP**GV(Nle)GFO**GP***x*** where the *x*-position was varied to Pro (pep-P, capable of recognizing both DDRs), Lys (pep-K) or Asp (pep-D) (**supplementary table 1**). All three THPs were flanked by GPP-rich domains to promote triple helix formation and a C-terminal Tyr for accurate spectrophotometric determination of concentration. The N-terminal flanking triplet of GVMGFO in the known DDR-binding sites varies between GQP (Col2 and Col3),^49,50^ GQA (*α*1 chain of Col1) and GEP (*α*2 chain of Col1)^64^. We introduced GPP in this position of the THPs to minimize unintended bias in the recognition specificity due to this variation. The rationale for introducing Lys/Asp in the *x-*position of GP**R**GPP**GV(Nle)GFO**GP***x*** arises from the potential DDR2-binding motif GPIGPP**GIPGFP**GS***K*** found in Col10, which contains a Lys in equivalent sequence position. In the DDR2-THP crystal structure, the *x*-position residue is located close to H172 and the equivalently located R172 in DDR2.^52^ As discussed previously, H172R is also one of the five mutations that reverse the collagen-binding specificity of DDR2.^49^

The equilibrium population of triple-helical assemblies is sensitive to sequence variation and experimental conditions. To ensure that these factors do not confound the recognition specificities, we covalently captured all three THPs via Lys – Glu isopeptide bonds, following a protocol previously established by us^64^ (**see methods**). We introduced KGE sequence triplets on either side of the DDR-binding motif as shown in **supplementary table 1.** The KGE triplets are known to form salt bridges between the leading-middle and middle-trailing chain^73^. The carboxylate and ammonium head groups of the interacting residues in the salt bridges can be linked via an isopeptide bond using amide coupling. This covalently links the three peptides to each other, increasing stability and rendering the triple helix population independent of concentration. The isopeptide bonds are separated from the binding motif using inert GPP spacers, precluding unintended interaction between the isopeptide and the DDR DS-domains.

The covalently captured THPs are referred to as pep-P-cc, pep-K-cc and pep-D-cc, indicating the identity of the *x*-residue in the DDR binding motif GP**R**GPP**GVMGFO**GP***x*.** Following covalent capture, the peptides were purified by size-exclusion chromatography (SEC) to isolate the trimeric peptide species from residual monomers and reaction by-products (**supplementary fig. 4**). MALDI– TOF mass spectrometry was used throughout peptide synthesis, purification and covalent capture to track the molecular species. The uncaptured peptides displayed the expected monomeric masses (∼4.1 kDa), whereas the covalently captured peptides showed a predominant species at ∼ 12 kDa, consistent with the formation of triple-helical trimers (**supplementary fig. 5**). Circular dichroism (CD) spectroscopy was used to compare the stability of the monomeric and covalently captured peptides. Polypeptides within a collagen triple helix adopt a polyproline type II conformation, which results in a CD spectrum composed of a maximum between 220-225 nm and a minimum between 190-210 nm. Incidentally, unfolded monomer has a similar spectral profile but with a key difference – the maximum at ∼225 nm has a large positive ellipticity for a triple helix while that of the monomer is negative.^74^ Therefore, the difference in ellipticity at ∼225nm can be used to distinguish folded trimer from unfolded monomers. As shown in **supplementary figs. 6**, the CD spectrographs of non-covalently self-assembled peptides pep-P and pep-K at 10 °C show a maximum at ∼ 225 nm with a large positive intensity, suggesting stable triple helices. Upon heating to 37 °C, the ellipticity at 225 nm decreases to below zero, indicating unfolding of the trimer. However, the spectrum of pep-D at 10 °C shows a weakly positive maximum at 225 nm, suggesting a comparatively smaller population of triple helices and likely lower stability then pep-P or pep-K. Covalent capture increases the stability all three THPs as reflected in a large increase in normalized ellipticity (referred to as molar residue weight, abbreviated MRW) at 10 °C as compared to native THPs and relatively smaller decrease in MRW upon heating to 37 °C. The increased stability upon covalent capture is also reflected in higher thermal unfolding temperature of pep-P-cc and pep-K-cc, as compared to pep-P and pep-K, respectively (**supplementary fig. 7**). Crucially, while we do not observe an unfolded transition for pep-D, suggesting that this peptide does not form a triple helix, its covalently captured counterpart shows a cooperative unfolding transition with melting temperature of 41 °C. In the case of DDRs, binding of the collagen ligand activates the receptor. This activation assay has previously been performed by incubating THPs with cell-expressed DDRs at 37 °C for > 90 minutes.^48^ As shown in **supplementary fig. 8**, covalent capture protects against unfolding under this prolonged heating conditions: Pep-P and pep-K unfold to monomers in less than 30 minutes whereas pep-D remains unfolded. Upon covalent capture, the ellipticities of all three THPs show small decrease in the first few minutes followed by a stable positive ellipticity over at least 80 minutes. These results emphasize the importance of covalently capturing THP constructs, especially during comparative assessment of their affinities to other proteins and activation assays.

### Electrostatically charged residues influence THP specificities for DDR1 and DDR2

We first assessed the binding of pep-P-cc, pep-K-cc and pep-D-cc to Fc-linked DS-domains of DDR1 and DDR2 using static solid phase ELISA (**see methods**). As shown in **Fig. 3A**, all three THPs bind DDR1 and DDR2 but the endpoint optical density for DDR1 is less than that of DDR2 for pep-K-cc and pep-D-cc, suggesting lower binding affinity. This difference is confirmed by dose-response curves shown in **Fig. 3B-D**.

**Fig. 3.**
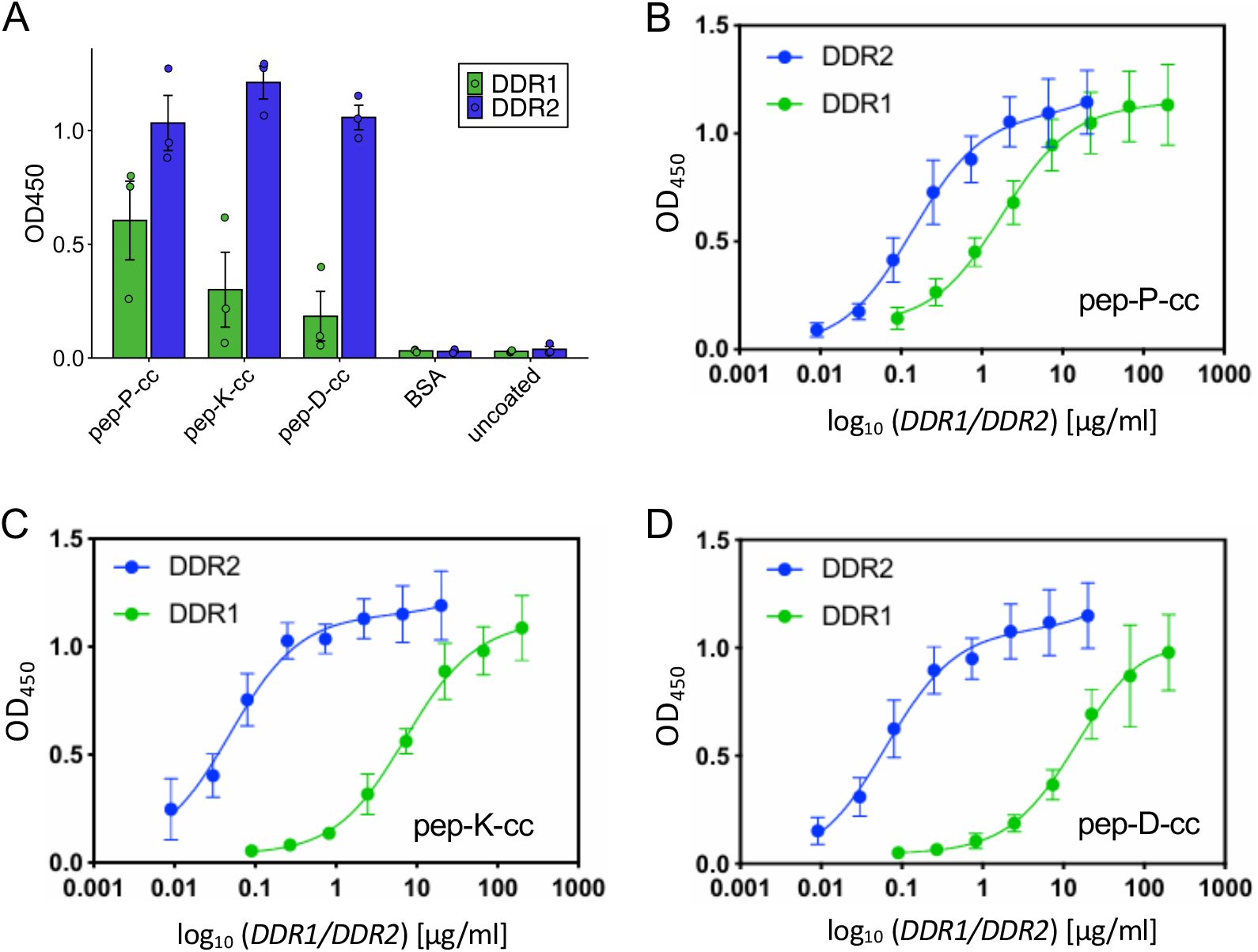
Analysis of DDR Binding to THPs assessed with ELISA. (**A**) Static ELISA binding assay of the covalently captured peptides pep-P-cc, pep-K-cc and pep-D-cc. Plates were coated with peptides at 1 µg/mL. DDR1-Fc and DDR2-Fc were applied at concentrations of 20 µg/mL and 2.5 µg/mL, respectively. BSA and uncoated wells served as negative controls. Data represent the mean optical density at 450 nm (OD_450_). Error bars represent standard deviation (SD) of the fit from triplicate measurements performed in two independent experiments. (**B–D**) Dose-response ELISA binding curves for pep-P-cc, pep-K-cc and pep-D-cc. Plates were coated with peptide at 1 µg/mL. DDR1-Fc and DDR2-Fc were applied as eight-point 1:3 serial dilutions starting from 200 µg/mL (DDR1) and 20 µg/mL (DDR2), respectively. Data represent the mean OD_450_ ± SD from technical duplicates obtained in two independent experiments. Curves represent the nonlinear regression fit of the binding data to a single-site binding model using GraphPad Prism.

All three THPs have a higher affinity for DDR2 but specificities, judged by the ratio of their dissociation constants, vary significantly with the nature of the electrostatic charge at the *x*-position of GP**R**GPP**GVMGFO**GP***x*** (**Table 2**). The ratio of dissociation constant (DDR1/DDR2) for pep-P-cc is 18 while that for pep-K-cc is 144. Substituting Lys with Asp further increases the ratio to 255. It is interesting to note that the nature of the *x*-position residue does not significantly affect binding affinity to DDR2 when the confidence interval of the dissociation constants is taken into consideration. In contrast, DDR1 affinity for the THPs is more sensitive to charge, where an 8 and 4-fold decrease in affinity is observed for the Asp and Lys-containing sequences, respectively. The lowering of DDR1 binding affinity to pep-K-cc and pep-D-cc primarily drives the increased specificity of THPs for the DDR2. These results indicate that electrostatic interactions are more important for recognition specificity of DDR1 than DDR2, supported also by the requirement of the N-term Arg flanking the consensus motif to bind DDR1^50^. Thus, there are two sequence positions that appear to determine DDR1-binding specificity: at Arg N-terminal to the minimal recognition motif and at Lys C-terminal to it. As discussed in the previous sections, the DS-domains of DDR1 and DDR2 differ substantially in the distribution and density of electrostatic charges close to these two ligand sequence positions. It is reasonable to extrapolate that electrostatic interactions in these two regions guide the recognition specificity.

**Table 2:** Dissociation constants for the binding of THPs to DDR1 and DDR2.

| THP | $K_{d,DDR1}$ (95% CI*)<br>ug ml<sup>-1</sup> | $K_{d,DDR2}$ (95% CI*)<br>ug ml<sup>-1</sup> | $K_{d,DDR1}/K_{d,DDR2}$ |
| --- | --- | --- | --- |
| pep-P-cc | 1.8 (1.1-3.0) | 0.1 (0.1-0.2) | 18 |
| pep-K-cc | 7.2 (4.7-10.8) | 0.05 (0.03-0.08) | 144 |
| pep-D-cc | 15.3 (7.8-31.1) | 0.06 (0.04-0.1) | 255 |

\*confidence interval

## Discussion

DDR1 and DDR2 tyrosine kinases play a central role in regulating cell migration and tissue remodelling during physiological development.^75^ Dysfunction in either receptor contributes to development of cardiovascular diseases^13^, fibrosis^12^ and several forms of cancers^16^. While several aspects of DDR biology related to its activation and subsequent regulation of intracellular signalling and cell response have been revealed, the molecular basis of its interaction with the different collagen types has not been fully elucidated. In a landmark work, Hohenester and co-workers determined the crystal structure of DDR2 DS-domain bound to a THP carrying the consensus motif conserved in Col2 and Col3, which revealed molecular determinants of this interaction and also provided broad contours of how collagens might interact with DDR1.^52^ We recently employed computationally designed heterotrimeric THPs to show that a similar motif found in the Col1 (*α*1)_2_*α*2 heterotrimer also recognizes both DDRs.^64^ However, the two DDRs together recognize an additional four types of collagens Col4, Col5, Col6 and Col10 with varying specificity.^5,6^ We set out to investigate the molecular basis of this collagen-binding specificity of DDRs. Our results with THPs combined with observations from biochemical data published on collagen-DDR interaction suggests an important role for electrostatic interactions in determining specificity.

Proteins often employ electrostatic interactions to improve ligand specificity in a competitive setting. A classic comparison is chymotrypsin vs trypsin, which are structurally related serine proteases with different preferences for substrate residues. Trypsin contains an Asp positioned in its specificity pocket that favours substrates with basic Arg/Lys residues.^76^ Substitution of Asp for Ser, found in chymotrypsin, causes five orders of magnitude decrease in selectivity for such substrates.^77^ G-protein coupled receptors (GPCRs) present another important example. Human bradykinin B1 and B2 are related GPCRs with different selectivity for bradykinin-like peptide ligand – B2 prefers peptides with a C-terminal Arg whereas B1 prefers peptides lacking the Arg. B1 receptor contains a strategically placed Lys, occupied by Ser in B2, that likely dictates ligand selectivity via electrostatic forces.^78^ In case of DDRs, the primary ligand binding force has been understood to be the hydrophobic interaction between the ligand phenylalanine and the amphipathic binding pocket on the DS-domains.^52^ However, the observation that the absence of the N-terminal Arg abrogates binding of consensus motif GVMGFO to DDR1^43^ suggest a predominant role of electrostatic interactions in determining not only specificity but any binding in the first place. Further, the placement of Lys and Asp on the C-terminal of the consensus motif in our THPs significantly influences the THP affinity for DDR1. This ligand sequence position is located within interacting distance of R172 in DDR1 and H172 in DDR2 while also being in the vicinity of the acidic patch on the DDR2 DS-domains. Thus, it appears that the electrostatic interactions between charged residues towards the N-and C-terminal of the minimal motif GVMGFO, the acidic patches of the DDR1 and DDR2 and the residue in sequence position 172 determines the recognition specificity.

Free energy change due to salt bridge interactions is determined by a combination of Coulombic forces, desolvation penalty of the charged head group of the interacting residues and conformational entropic loss of the sidechains.^79^ Thus, despite the simplistic repulsive/attractive mechanisms suggested by examples above, electrostatic interactions in proteins are sophisticated and fine-tuned lever to modulate ligand thermodynamics and kinetics. For example, electrostatic interactions modulate binding kinetics i.e. control how fast the ligand binds to the target (association rate, *K_on_)* and how quickly it comes off (dissociation rates, *K_off_*). Due to their long-range nature, electrostatic interactions typically increase association rates by guiding binding partners together faster than simple diffusion, called electrostatic steering^80^, and also by stabilizing the transiently formed states before final stereospecific complexation occurs.^81^ The strong ionic bonds also act as an electrostatic trap to slow the rate of ligand dissociation.^82^ Inverse of the off-rate, called the ligand residence time, is the amount of time a ligand stays bound to the target. Going beyond thermodynamics of interactions, recent attention in drug targeting has pivoted to modulating the ligand residence time, which can determine how long the therapeutic effect lasts in a living system.^83^ Longer residence time can also improve specificity of a ligand for one target over the other, even if both have similar affinities. This kinetic trapping effect could be relevant to DDRs as well. It is known that upon ligand binding, DDRs undergo slow and sustained phosphorylation, often taking hours to peak and then detectable for days.^5,6^ In addition, it has also been hypothesized that collagen facilitates lateral oligomerization of DDR1, which is a pre-requisite for phosphorylation.^84^ It can be extrapolated that the ligand remains bound for the duration of this event, suggesting long residence times. DDR2 recognizes several toolkit THPs but only those containing the consensus motif and the N-terminal Arg are able to activate the receptor and cause intracellular phosphorylation – truncated sequences lacking the N-terminal Arg bind but not activate the receptor.^50^ It could be hypothesized that this Arg is required to achieve sufficiently long residence time for receptor oligomerization and activation. In the case of DDR1, Arg is critical for binding. Since the collagen-binding pocket of DDR1 is less negatively charged than that of DDR2, it can also be hypothesized that ligand residence time of the Arg containing ligand would be shorter than DDR2. The requirement for long residence time likely also prevents other similar motifs, shown to bind DDRs in the THP toolkit experiments ^49,50^, from activating the receptor perhaps because they are not bound long enough. Our future work will focus on comparing the on-and off-rates for the interaction of the ligand with both DDRs and also dissecting the effect of the N-terminal Arg on these rates. Further, we will design new THPs to systematically test how charged residues in sequence positions on either side of the minimal motif GVMGFO not tested in this study influence equilibrium affinity and binding kinetics.

Therapeutic agents employed in treatment of DDR-related pathologies and especially cancer either target its intracellular kinase activity or the ectodomains responsible for collagen recognition.^85^ The kinase domains are structurally homologous across several RTKs^86^ including DDRs^75^. Thus, small molecule drugs, frequently used as inhibitors of kinase activity, indiscriminately target kinases in other RTKs, causing unwanted side effects.^87^ Thus, therapeutic agents that inhibit the collagen-DDR interaction are more attractive.^88^ Further, given their highly differentiated function, it is often desirable to specifically target either DDR1 or DDR2.^16,75^ Molecular understanding of how electrostatic interactions determine DDR recognition specificity could contribute to development of specific inhibitors of collagen-DDR interactions.

## Material and methods

### Isoelectric point analysis

The isoelectric point of the DS-domains was calculated on the webserver https://www.protpi.ch/Calculator/ProteinTool using corresponding sequences obtained from UniProt^89^ (human DDR1, id Q08345, residues 31-185; human DDR2, id Q16832, residues 30-185). The titration curve reported by the webserver was used to obtain the net charge at physiological pH of 7.4 (**supplementary fig. 1**).

### Surface electrostatic potential analysis

The surface electrostatic potentials were calculated using the Adaptive Poisson-Boltzmann Solver (APBS)^90^ electrostatics plugin in PyMOL^91^. Previously published crystal structures of the DDR2 DS-domain with a collagen peptide (pdb:2wuh^52^) and the extracellular domain of DDR1 with an inhibitory Fab fragment^92^ were trimmed to the DS-domains (residues 31-185 for DDR1 and residues 30-185 for DDR2). These input structures were prepared by PDB2QR^93,94^ to build missing atoms, add and optimize hydrogens and calculate protonation states based on residue pKa values. The prepared structures were then fed into the APBS^90^ for calculation of surface electrostatic charge profile and rendered as surfaces coloured from −5 kbTec −1 (red) to +5 kbTec −1 (blue) in PyMOL^91^.

### Solid phase peptide synthesis

Peptides were synthesized on a Liberty Blue 2.0 Microwave Peptide Synthesizer (CEM, Kamp Lintfort, Germany) following solid-phase peptide synthesis (SPPS) with cycles of orthogonal Fmoc (9-fluorenylmethoxycarbonyl) based coupling chemistry, essentially as reported previously.^64^ In a typical cycle, 1.0 M solutions of amino acids were coupled onto a polystyrene-based TentaGel S RAM resin (Rapp Polymere GmbH; loading = 0.24 mmol g^-1^) using cycles of coupling with 1.0 M OxymaPure® (ethyl-2-cyano-2-(hydroxyamino)acetate) and 0.5 M N,N’-diisopropylcarbodiimide (DIC) and Fmoc-deprotection with 10 % v/v piperidine. Solutions of all reagents were prepared in N, N-dimethylformamide (DMF). Prior to coupling, resins were swollen in DMF and transferred to the reaction chamber. Coupling of aspartic acid was performed with a bulky O-3-methyl-pent-3-yl (OMpe) group sidechain protection to prevent aspartimide formation via intramolecular cyclization. Similarly, to minimize δ-lactam formation and subsequent residue deletion, arginine coupling cycles were performed twice at room temperature in absence of microwave heating. Further, to improve coupling efficiency and minimize deletion, we double coupled all instances of tandem prolines. After coupling of the final N-terminal amino acid, Fmoc was removed via two cycles of deprotection and resins transferred to a 10 ml polypropylene syringe fitted with a silica frit. The resins were washed five times with 5 ml each of dichloromethane (DCM) and dried for a minimum of 30 minutes under ambient conditions. The dry resins were N-terminally acetylated using a mixture of 250 µL N-ethyldiisopropylamine (DIEA), and 250 µL acetic anhydride ((Ac)₂O) in 20 mL DCM. Following acetylation, the resins were washed again with DCM and dried. For cleavage of the peptide from the resin, a cleavage cocktail comprising 250 µL 2,2’-(ethylenedioxy)diethanethiol (DODT), 250 µL triisopropylsilane (TIS), 250 µL deionized water (H₂O MQ) 9.25 mL trifluoroacetic acid (TFA) was used. The cleaved peptides were precipitated by slow dripping of the cleavage solution into ice-cold (−20°C) diethyl ether. The precipitates were collected via vacuum filtration using a PTFE filter (0.45 µm, Sartorius Stedim Biotech GmbH) and washed five times with 5 ml diethyl ether each. The peptides were then allowed to dry under ambient conditions and stored at −20°C.

### Covalent capture of triple helices

The triple-helical peptides were covalently captured using a method previously established by Jalan et al.^64^ Briefly, each peptide contained a KGE motif capable of forming three interchain lysine – glutamate salt bridges in a triple helix. Capture was achieved by covalently linking the lysine and glutamate head groups within the salt bridge via an isopeptide bond. In a typical protocol, stock solutions of peptides (3 mM), EDC (600 mM) and OxymaPure (36 mM) were prepared in 100 mM MES buffer at pH 6.8 and stored in cold room maintained at 5 °C. The original protocol employed 4-hydroxybenztriazole (HOBt) but, due to its explosive nature, we substituted it with OxymaPure. The stock solutions were mixed to obtain a stoichiometric ratio of 1:40:4 for peptide, EDC and OxymaPure, respectively, vortexed and stirred in cold room. To monitor the progress of the covalent capture reaction, aliquots from the reaction mixture were quenched with 0.2 M hydroxylamine and analysed using HPLC and mass spectrometry. Upon completion of the reaction as assessed by HPLC chromatogram profiles, the entire reaction mixture was quenched with equal volume of 0.2 M hydroxyl and desalted using centrifugal filters with a 2 kDa molecular weight cut-off (Vivaspin® 15R Centrifugal Concentrator Hydrosart®, Sartorius AG, United Kingdom). These concentrators also facilitated buffer exchange from the MES buffer used in the covalent capture reaction to 1x Dulbecco’s PBS (DPBS) needed for purification via size-exclusion chromatography (SEC) and ELISA assays. Buffer exchange was achieved by washing the samples five times with 1× PBS.

### Chromatographic purification of peptides

Crude monomeric peptides were purified via high performance liquid chromatography (HPLC) on a Phenomenex semi-preparative column (Aeris peptide XB-C18 column (Aeris^TM^ 5µm PEPTIDE XB-C18 100 column, LC Column 250 x 10.0 mm). Method for the semi-prep purification were optimized with on Phenomenex analytical column (Aeris^TM^ 5µm PEPTIDE XB-C18 100, LC Column 250 x 4.6 mm). For both columns, the mobile and elution phases were 0.1% aqueous trifluoroacetic acid (TFA) and 97.5% aqueous acetonitrile containing 0.085% TFA. Elution peaks were monitored with a JASCO UV-2075 Plus UV-detector using JASCO ChromNAV version 2.04.05 for method modification, execution, and data analysis. In a typical HPLC run, the samples were dissolved at a concentration of 10-20 mg/ml in 50 % aqueous acetonitrile, centrifuged at 13,300 rpm for 10 minutes at room temperature, the supernatant collected and loaded onto a 100 μl loading loop. Upon injection, a gradient ranging from 0.5 to 1 % aqueous acetonitrile at a flow rate of 6 ml min^-1^ was run. 0.5 to 1 ml fractions were collected at the main peak and analysed by mass spectrometry. Fractions displaying peaks corresponding to the desired mass were pooled, acetonitrile removed via rotary evaporation at room temperature, flash frozen in liquid nitrogen and lyophilized to obtain the dry purified peptide powder used for all subsequent analyses.

### Desalting, concentration and Buffer exchange

To monitor the progress of the covalent capture reaction, aliquots were collected and analysed using HPLC and mass spectrometry. Prior to these analyses, the aliquots underwent desalting using C18 Zip-Tip® (ZIPTIP MIT 0.6 UL C18-HARZ, Sigma-Aldrich Chemie GmbH, Germany). The Zip-Tips were first activated with 100% acetonitrile (AcN) and subsequently washed with a solution of H₂O containing 0.1% trifluoroacetic acid (TFA). The peptide samples were then loaded onto the tips by carefully aspirating the solution 7–10 times, ensuring efficient binding of the peptides to the filter. Any residual salts were removed with an additional H₂O + 0.1% TFA wash, and the peptides were eluted in 60% AcN for further processing. For larger sample volumes, desalting was performed via dialysis using 100–500 Da dialysis membranes (Spectra/Por® Dialysis Membrane Biotech CE Tubing, Spectrum Laboratories Inc., Rancho Dominguez, USA) against Milli-Q water (H₂O MQ).

To concentrate the peptide solutions, either lyophilization or centrifugal concentration was employed. Centrifugal concentrators equipped with 2 kDa or 5 kDa membranes (Vivaspin® 15R Centrifugal Concentrator Hydrosart®, Sartorius AG, United Kingdom) were used to reduce sample volume while preserving peptide integrity. These concentrators also facilitated buffer exchange from the MES buffer used in the covalent capture reaction to 1× PBS, which was required for subsequent size-exclusion chromatography (SEC) and ELISA assays. Buffer exchange was achieved by washing the samples five times with 1× PBS.

### Mass spectrometric characterization of peptides and covalently captured triple helices

Masses of crude, HPLC purified and covalently captured peptides were determined using autoflex® maX (Bruker Daltonics GmbH & Co. KG) Matrix-assisted Laster Desorption/Ionization Spectroscopy – Time of Flight (MALDI-TOF) spectrometer. Crude peptides were dissolved in 50% aqueous acetonitrile and the resulting solution diluted 1:1 in 60% w/v matrix solution of α-Cyano-4-hydroxycinnamic (HCCA) in acetonitrile. 3 μL of the peptide was spotted onto the MALDI plate and dried under ambient conditions before mass spectrometric analysis. Fractions of HPLC purified monomeric peptides were directly mixed in 1:1 v/v ratio with the HCCA matrix solution and spotted onto the plate. Aliquots of the covalently captured peptides were desalted using C18 Zip-Tip® (ZIPTIP MIT 0.6 UL C18-HARZ, Sigma-Aldrich Chemie GmbH, Germany). The Zip-Tips were first activated with 100% acetonitrile and subsequently washed with a solution of H₂O containing 0.1% TFA. The peptide samples were then loaded onto the tips by carefully aspirating the solution 7–10 times, ensuring efficient binding of the peptides to the filter. Any residual salts were removed with an additional H₂O + 0.1% TFA wash, and the peptides were eluted in 60% acetonitrile. 1:1 v/v mixture of the eluted solution in HCCA matrix was spotted onto the MALDI plate for further analysis. In all cases, the MALDI-TOF mass spectra were baseline corrected and exported as text files using Bruker AutoFlex Analysis software. Data analysis and peak interpretation were carried out using the exported text files in mMass version 5.5.0.^95^

### Size-Exclusion Chromatography (SEC)

To separate monomeric peptides from covalently bound triple-helical peptides and any potential multimers, size-exclusion chromatography (SEC) was performed using a Superdex 75 10/300 GL column (Sigma-Aldrich Chemie GmbH, Germany) running an isocratic gradient of 1x DPBS at 0.8 ml per min. Prior to SEC, the samples were concentrated to a final loading volume of 200 µL using a 5 kDa Vivaspin centrifugal concentrator. Fractions collected at 1 ml interval were analysed by mass spectrometry and those with the desired trimer were pooled, concentrated and the final triple-helical concentration determined using tyrosine extinction coefficient of 1399 M^-1^ cm^-1^ at 275 nm on a Nanodrop ND-1000 spectrophotometer (Thermo Fisher Scientific, Wilmington, DE, USA) for further use in solid-phase binding assays with DDRs.

### Evaluation of DDR binding using Enzyme-Linked Immunosorbent Assay (ELISA)

Binding of DDR1 and DDR2 to the covalently captured peptides was assessed by a combination of static single concentration and dose response indirect ELISA. For static ELISA, 96-well plates (Immulon 2HB, Thermo Fisher Scientific) were coated with peptides at a concentration of 10 µg/mL in 1× PBS buffer overnight at 4°C. After incubation, the peptide solutions in the wells were discarded and the surface blocked with 200 µL of 2% bovine serum albumin (BSA, Fraktion V, Carl ROTH GmbH, Germany) to minimize non-specific binding. Following three washes with a washing buffer containing 0.1% BSA and 0.05% Tween^®^20 (Carl ROTH GmbH, Germany), the coated wells were incubated with either 100 µL of 250 ng/mL DDR1 or 100 ng/mL DDR2 in 1× PBS for 2 hours at room temperature. DDR1 consisted of the extracellular region of DDR1 fused to a human IgG Fc fragment, while DDR2-Fc comprised the collagen-binding discoidin homology domain of DDR2 fused to a human IgG Fc fragment.^49^ After incubation, the wells were washed three times with the washing buffer and then incubated for 1 hour at room temperature with 100 µL of horseradish peroxidase (HRP)-conjugated anti-human Fc antibody (0.8 mg/mL, Jackson ImmunoResearch, West Grove, PA, USA), diluted 1:10,000 in PBS. Unbound antibodies were thoroughly removed by washing the wells five times with washing buffer. Binding was detected using the 1-Step™ Ultra TMB-ELISA Substrate Solution (Thermo Fisher Scientific, USA) that facilitates the catalytic conversion of 3,3ʹ,5,5ʹ-tetramethylbenzidine (TMB) and hydrogen peroxide (H₂O₂) with HRP to an oxidized product with blue colour. After development of a blue colour in the plate, the reaction was quenched with 50 µL of 2N sulfuric acid (H₂SO₄), resulting in a colour change from blue to yellow. Absorbance was measured at 450 nm using a SpectraMax iD5 plate reader (Molecular Devices, San Jose, USA). Dose response curves were obtained with plates coated with 200 µl of 1 µg/ml peptide solutions incubated with 1:3 serial dilutions of DDR1 and DDR2. Dissociation constants were calculated using GraphPad Prism (Version 10.5.0, ©1992-2025 GraphPad Software, LLC).

### Assessment of the peptide thermal stability using circular dichroism (CD)

The secondary structure of the purified monomeric peptides and covalently captured triple helices in solution and their thermal stabilities were determined using CD spectroscopy. In a typical experiment, stock solution of peptides prepared in miliQ water were diluted to obtain a final concentration of 0.5 to 1 mg/ml in 10 mM sodium phosphate buffer at pH 7.4. The stock solutions of covalently captured peptides in DPBS were similarly diluted to obtain 1 mg/ml solutions but in 0.1 X DPBS. The solutions were transferred to a quartz cuvette with 1mm pathlength and analysed on a JASCO 810 spectropolarimeter. The secondary structures were determined by scanning from 250 to 190 nm at rate of 100 nm min^-1^, data pitch of 1 nm and a response time of 4 s. The CD spectrum scan were performed both at 10°C and 37°C to assess the change in secondary structure. The thermal stability of the peptides and covalently captured constructs were determined via variable temperature measurements and isothermal unfolding. In a variable temperature measurement, the change in the CD ellipticity signal at 225 nm was monitored between 15 and 75 °C while heating at the rate of 0.2 °C min^-1^. Isothermal unfolding scans were obtained by monitoring ellipticity at 225 nm as a function of time at the constant temperature of 37 °C. Data from the wavelength scan and thermal and isothermal unfolding were exported as text files and plotted in GraphPad Prism version 10.5.0.

## Author’s contributions

AAJ conceived the research idea, designed the experiments, analysed and interpreted the data with MP and LJ and co-wrote the manuscript with MP, LJ and BL. LJ prepared the initial sets of peptides to test the DDR-binding specificity and performed the initial ELISA experiments, concluding that covalent capture is necessary to fully reveal the recognition specificity. MP refined these initial experiments with additional peptides, performed covalent capture, obtained static and dose-response ELISA data and associated CD data, analysed and plotted the data and with LJ contributed to the revision of the manuscript. JDM prepared the initial set of peptides and contributed to the revision of the manuscript. BL provided the DDR-Fc constructs, performed experiments that precipitated the need to perform a covalent capture of peptides and contributed to research design and revision of the manuscript.

## Declaration of generative AI and AI-assisted technologies in the writing process

The authors did not use generative large-language model-based tools to write any part of this manuscript.

## Acknowledgements

Funding was provided by the funding scheme Exzellenzverbünde und Universitäts-kooperationen (EVUK) within the project “Function by Design: Cellular Hybrids” by the Bavarian Ministry of the Science and Art. LJ is funded by the Elite Network of Bavaria as a member of the Elite Graduate Program “Biological Physics”. The authors are thankful to Vamika Sagar and Mateo Ceballos Giraldo for support during peptide synthesis and purification.

## Supplementary materials

Supplementary material associated with article is supplied along with the manuscript and can be found online at hyperlink.

## Data availability

Data underlying this manuscript can be requested from the corresponding author upon reasonable request.

