## Supplementary Material for "Molecular Insights into the Collagen-binding Specificity of Discoidin Domain Receptors"

*† equal contribution*

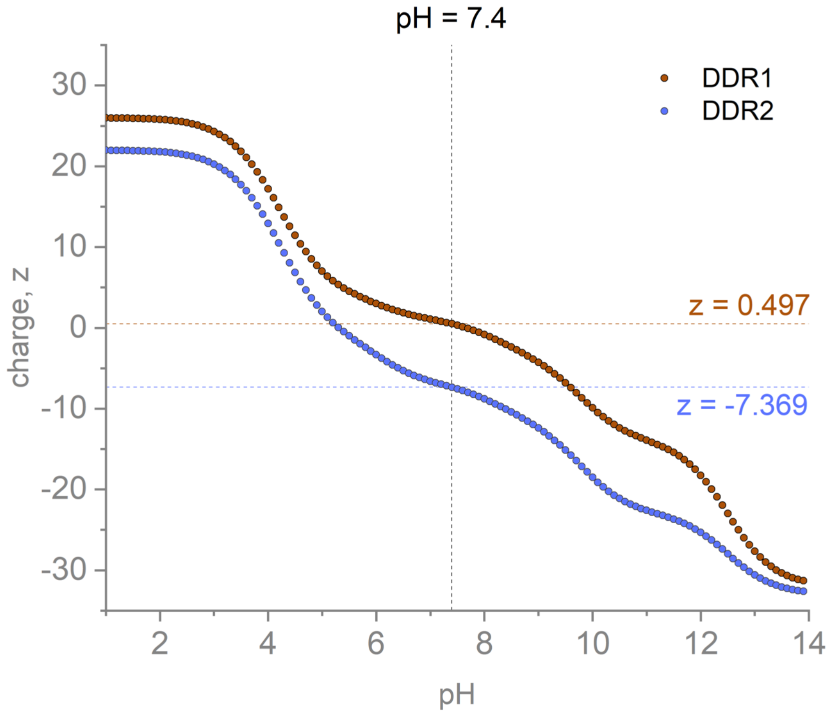


**Supplementary Fig. 1.** Isoelectric point of the DDR DS-domains as calculated on the webserver https://protpi.ch using default settings. Sequences of the DS-domains of human DDR1 (entry id: Q08345, residues 31-185) and DDR2 (entry id: Q16832, residues 30-185) were obtained from UniProt[1] knowledge database (UniProtKB).

### Col10 - $(\alpha 1)_3$

```
α1(X)    1 MLPQIPFLLLVSLNLVHGVFYAERYQMPTGIKGPLPNTKTQFFIPYTIKSKGIAVRGEQGTPGPPGPAGPRGHPGPSGPPGKPGYGSPGLQGEPGLPGPP 100
α1(X)    1 MLPQIPFLLLVSLNLVHGVFYAERYQMPTGIKGPLPNTKTQFFIPYTIKSKGIAVRGEQGTPGPPGPAGPRGHPGPSGPPGKPGYGSPGLQGEPGLPGPP 100
α1(X)    1 MLPQIPFLLLVSLNLVHGVFYAERYQMPTGIKGPLPNTKTQFFIPYTIKSKGIAVRGEQGTPGPPGPAGPRGHPGPSGPPGKPGYGSPGLQGEPGLPGPP 100

α1(X)  101 GPSAVGKPGVPGLPGKPGERGPYGPKGDVGPAGLPGPRGPPGPPGIPGPAGISVPGKPGQQGPTGAPGPRGFPGEKGAPGVPGMNGQKGEMGYGAPGRPG 200
α1(X)  101 GPSAVGKPGVPGLPGKPGERGPYGPKGDVGPAGLPGPRGPPGPPGIPGPAGISVPGKPGQQGPTGAPGPRGFPGEKGAPGVPGMNGQKGEMGYGAPGRPG 200
α1(X)  101 GPSAVGKPGVPGLPGKPGERGPYGPKGDVGPAGLPGPRGPPGPPGIPGPAGISVPGKPGQQGPTGAPGPRGFPGEKGAPGVPGMNGQKGEMGYGAPGRPG 200

α1(X)  201 ERGLPGPQGPTGPSGPPGVGKRGENGVPGQPGIKGDRGFPGEMGPIGPPGPQGPPGERGPEGIGKPGAAGAPGQPGIPGTKGLPGAPGIAGPPGPPGFGK 300
α1(X)  201 ERGLPGPQGPTGPSGPPGVGKRGENGVPGQPGIKGDRGFPGEMGPIGPPGPQGPPGERGPEGIGKPGAAGAPGQPGIPGTKGLPGAPGIAGPPGPPGFGK 300
α1(X)  201 ERGLPGPQGPTGPSGPPGVGKRGENGVPGQPGIKGDRGFPGEMGPIGPPGPQGPPGERGPEGIGKPGAAGAPGQPGIPGTKGLPGAPGIAGPPGPPGFGK 300

α1(X)  301 PGLPGLKGERGPAGLPGGPGAKGEQGPAGLPGKPGLTGPPGNMGPQGPKGIPGSHGLPGPKGETGPAGPAGYPGAKGERGSPGSDGKPGYPGKPGLDGPK 400
α1(X)  301 PGLPGLKGERGPAGLPGGPGAKGEQGPAGLPGKPGLTGPPGNMGPQGPKGIPGSHGLPGPKGETGPAGPAGYPGAKGERGSPGSDGKPGYPGKPGLDGPK 400
α1(X)  301 PGLPGLKGERGPAGLPGGPGAKGEQGPAGLPGKPGLTGPPGNMGPQGPKGIPGSHGLPGPKGETGPAGPAGYPGAKGERGSPGSDGKPGYPGKPGLDGPK 400

α1(X)  401 GNPGLPGPKGDPGVGGPPGLPGPVGPAGAKGMPGHNGEAGPRGAPGIPGTRGPIGPPGIPGFPGSKGDPGSPGPPGPAGIATKGLNGPTGPPGPPGPRGH 500
α1(X)  401 GNPGLPGPKGDPGVGGPPGLPGPVGPAGAKGMPGHNGEAGPRGAPGIPGTRGPIGPPGIPGFPGSKGDPGSPGPPGPAGIATKGLNGPTGPPGPPGPRGH 500
α1(X)  401 GNPGLPGPKGDPGVGGPPGLPGPVGPAGAKGMPGHNGEAGPRGAPGIPGTRGPIGPPGIPGFPGSKGDPGSPGPPGPAGIATKGLNGPTGPPGPPGPRGH 500

α1(X)  501 SGEPGLPGPPGPPGPPGQAVMPEGFIKAGQRPSLSGTPLVSANQGVTGMPVSAFTVILSKAYPAIGTPIPFDKILYNRQQHYDPRTGIFTCQIPGIYYFS 600
α1(X)  501 SGEPGLPGPPGPPGPPGQAVMPEGFIKAGQRPSLSGTPLVSANQGVTGMPVSAFTVILSKAYPAIGTPIPFDKILYNRQQHYDPRTGIFTCQIPGIYYFS 600
α1(X)  501 SGEPGLPGPPGPPGPPGQAVMPEGFIKAGQRPSLSGTPLVSANQGVTGMPVSAFTVILSKAYPAIGTPIPFDKILYNRQQHYDPRTGIFTCQIPGIYYFS 600

α1(X)  601 YHVHVKGTHVWVGLYKNGTPVMYTYDEYTKGYLDQASGSAIIDLTENDQVWLQLPNAESNGLYSSEYVHSSFSGFLVAPM                     680
α1(X)  601 YHVHVKGTHVWVGLYKNGTPVMYTYDEYTKGYLDQASGSAIIDLTENDQVWLQLPNAESNGLYSSEYVHSSFSGFLVAPM                     680
α1(X)  601 YHVHVKGTHVWVGLYKNGTPVMYTYDEYTKGYLDQASGSAIIDLTENDQVWLQLPNAESNGLYSSEYVHSSFSGFLVAPM                     680
```

Col4 - $(\alpha1)_2\alpha2$

```
α1(IV)    1 ---------MGPRLSVWLLLLPAALLLHEEHSRAAA--KGGCAGSGC-GKCDCHGVKGQKGERGLPGLQGVIGFPGMQGPEGPQGPPGQKGDTGEPGLPGTKGTRGPPGASG  100
α1(IV)    1 ---------MGPRLSVWLLLLPAALLLHEEHSRAAA--KGGCAGSGC-GKCDCHGVKGQKGERGLPGLQGVIGFPGMQGPEGPQGPPGQKGDTGEPGLPGTKGTRGPPGASG  100
α2(IV)    1 MGRDQRAVAGPALRRWLLLGTVTVGFLAQSVLAGVKKFDVPCGGRDCSGGCQCYPEKGGRGQPGPVGPQGYNGPPGLQGFPGLQGRKGDKGERGAPGVTGPKGDVGARGVSG  112

α1(IV)  101 YPGNPGLPGIPGQDGPPGPPGIPGCNGTKGERGPLGPPGLPGFAGNPGPPGLPGMKGDPGEILGHVPGMLLKGERGFPGIPGTPGPPGLPGLQGPVGPPGFTGPPGPPGPPG  212
α1(IV)  101 YPGNPGLPGIPGQDGPPGPPGIPGCNGTKGERGPLGPPGLPGFAGNPGPPGLPGMKGDPGEILGHVPGMLLKGERGFPGIPGTPGPPGLPGLQGPVGPPGFTGPPGPPGPPG  212
α2(IV)  113 FPGADGIPGHPGQGGPRGRPGYDGCNGTQGDSGPQGPPGSEGFTGPPGPQGPKGQKGEP-YALPKEERDRYRGEPGEPGLVGFQGPPGRPGHVGQMGPVGAPGRPGPPGPPG  223

α1(IV)  213 PPGEKGQMGLSFQGPKGDKGDQGVSGPPGVPGQA-------QVQEKGDFATKGEKGQKGEPGFQGMPGVGEKGEPGKPGPRGKPGKDGDKGEKGSPGFPGEPGYPGLIGRQG  317
α1(IV)  213 PPGEKGQMGLSFQGPKGDKGDQGVSGPPGVPGQA-------QVQEKGDFATKGEKGQKGEPGFQGMPGVGEKGEPGKPGPRGKPGKDGDKGEKGSPGFPGEPGYPGLIGRQG  317
α2(IV)  224 PKGQQGNRGLGFYGVKGEKGDVGQPGPNGIPSDTLHPIIAPTGVTFHPDQYKGEKGSEGEPGIRGISLKGEEGIMGFPGLRGYPGLSGEKGSPGQKGSRGLDGYQGPDGPRG  335

α1(IV)  318 PQGEKGEAGPPGPPGIVIGTGPLGEKGERGYPGTPGPRGEPGPKGFPGLPGQPGPPGLPVPGQAGAPGFPGERGEKGDRGFPGTS-LP-GPSGRDGLPGPPGSPGPPGQPGY  427
α1(IV)  318 PQGEKGEAGPPGPPGIVIGTGPLGEKGERGYPGTPGPRGEPGPKGFPGLPGQPGPPGLPVPGQAGAPGFPGERGEKGDRGFPGTS-LP-GPSGRDGLPGPPGSPGPPGQPGY  427
α2(IV)  336 PKGEAGDPGPPGLP--AYSPHPSLAKGARGDPGFPGAQGEPGSQGEPGDPGLPGPPGLSIGDGDQRRGLPGEMGPKGFIGDPGIPALYGGPPGPDGKRGPPGPPGLPGPPGP  445

α1(IV)  428 TNGIVECQPGPPGDQGPPGIPGQPGFIGEIGEKGQKGESCLICDIDGYRGPPGPQGPPGEIGFPGQPGAKGDRGLPGRDGVAGVPGPQGTPGLIGQPGAKGEPGEFYFDLRL  539
α1(IV)  428 TNGIVECQPGPPGDQGPPGIPGQPGFIGEIGEKGQKGESCLICDIDGYRGPPGPQGPPGEIGFPGQPGAKGDRGLPGRDGVAGVPGPQGTPGLIGQPGAKGEPGEFYFDLRL  539
α2(IV)  446 DGFL-FGLKGAKGRAGFPGLPGSPGARGPKGWKGDAGECRCTEGDEAIKGLPGLPGPKGFAGINGEPGRKGDRGDPGQHGLPGFPGLKGVPGNIGAPGPKGAKGDS-RTITT  555

α1(IV)  540 KGDKGDPGFPGQPGMPGRAGSPGRDGHPGLPGPKGSPGSVGLKGERGPPGGVGFPGSRGDTGPPGPPGY---GPAGPIGDKGQAGFPGGPGSPGLPGPKGEPGKIVP-----  643
α1(IV)  540 KGDKGDPGFPGQPGMPGRAGSPGRDGHPGLPGPKGSPGSVGLKGERGPPGGVGFPGSRGDTGPPGPPGY---GPAGPIGDKGQAGFPGGPGSPGLPGPKGEPGKIVP-----  643
α2(IV)  556 KGERGQPGVPGVPGMKGDDGSPGRDGLDGFPGLPGPPGD-GIKGPPGDPGYPGIPGTKGTPGEMGPPGLGLPGLKGQRGFPGDAGLPGPPGFLGPPGPAGTPGQIDCDTDVK  666

α1(IV)  644 ---------------LPGPPGAEGLPGSPGFPGPQGDRGFPGTPGRPGLPGEKGAVGQPGI-GFPGPPGPKGVDGLPGDMGPPGTPGRPGFNGLPGNPGVQGQKGEP---GV  736
α1(IV)  644 ---------------LPGPPGAEGLPGSPGFPGPQGDRGFPGTPGRPGLPGEKGAVGQPGI-GFPGPPGPKGVDGLPGDMGPPGTPGRPGFNGLPGNPGVQGQKGEP---GV  736
α2(IV)  667 RAVGGDRQEAIQPGCIGGPKGLPGLPGPPGPTGAKGLRGIPGFAGADGGPGPRGLPGDAGREGFPGPPGFIGPRGSKGAVGLPGPDGSPGPIGLPGPDGPPGERGLPGEVLG  778

α1(IV)  737 GLPGLKGLPGLPGIPGTPGEKGSIGVPGVPGEHGAIGPPGLQGIRGEPGPPGLPGSVGSPGVPGI-GPPGARGPPGGQGPPGLSGPPGIKGEKGFPGFPGLD-MPGPKGDKG  846
α1(IV)  737 GLPGLKGLPGLPGIPGTPGEKGSIGVPGVPGEHGAIGPPGLQGIRGEPGPPGLPGSVGSPGVPGI-GPPGARGPPGGQGPPGLSGPPGIKGEKGFPGFPGLD-MPGPKGDKG  846
α2(IV)  779 AQPGPRGDAGVPGQPGLKGLPGDRGPPGFRGSQGMPGMPGLKGQPGLPGPSGQPGLYGPPGLHGFPGAPGQEGPLGLPGIPGREGLPGDRGDPGDTGAPGPVGMKGLSGDRG  890

α1(IV)  847 AQGLPGITGQSGLPGLPGQQGAPGIPGFPGSKGEMGVMGTPGQPGSPGPVGAPGLPGEKGDHGFPGSSGPRGDPGLKGDKGDVGLPGKPGSMDKVDMGSMKGQKGDQGEKGQ  958
α1(IV)  847 AQGLPGITGQSGLPGLPGQQGAPGIPGFPGSKGEMGVMGTPGQPGSPGPVGAPGLPGEKGDHGFPGSSGPRGDPGLKGDKGDVGLPGKPGSMDKVDMGSMKGQKGDQGEKGQ  958
α2(IV)  891 DAGFTGEQGHPGSPGFKGIDGMPGTPGLKGDRGSPGMDGFQGMPGLKGRPGFPGSKGEAGFFGIPGLKGLAGEPGFKGSRGDPGPPGPP-PVILPGMKDIKGEKGDEGPMGL 1001

α1(IV)  959 IGPIGEKGSRGDPGTPGVPGKDGQAGQPGQP-GPKGDPGISGTPGAPGLPGPKGSVGGMGLPGTPGEKGVPGIPGPQGSPGLPGDKGAKGEKGQAGPPGIGIPGLRGEKGDQ 1069
α1(IV)  959 IGPIGEKGSRGDPGTPGVPGKDGQAGQPGQP-GPKGDPGISGTPGAPGLPGPKGSVGGMGLPGTPGEKGVPGIPGPQGSPGLPGDKGAKGEKGQAGPPGIGIPGLRGEKGDQ 1069
α2(IV) 1002 KGYLGAKGIQGMPGIPGLSGIPGLPGRPGHIKGVKGDIGVPGIPGLPGFPGVAGPPGITGFPGFIGSRGDKGAPGRAGLYGEIGATGDFGDIGDT-INLPGRPGLKGERGTT 1112

α1(IV) 1070 GIAGFPGSPGEKGEKGSIGIPGMPGSPGLKGSPGSVGYPGSPGLPGEKGDKGLPGLDGIPGVKGEAGLPGTPGPTGPAGQKGEPGSDGIPGSAGEKGEPGLPGRGFPGFPGA 1181
α1(IV) 1070 GIAGFPGSPGEKGEKGSIGIPGMPGSPGLKGSPGSVGYPGSPGLPGEKGDKGLPGLDGIPGVKGEAGLPGTPGPTGPAGQKGEPGSDGIPGSAGEKGEPGLPGRGFPGFPGA 1181
α2(IV) 1113 GIPGLKGFFGEKGTEGDIGFPGITGVTGVQGPPGLKGQTGFPGLTGPPGSQGELGRIGLPGGKGDDGWPGAPGLPGFPGLRGIRGLHGLPGTKGFPGSPGSDIHGDPGFPGP 1224

α1(IV) 1182 KGDKGSKGEVGFP-GLAGSPGIPGSKGEQGFMGPPGPQGQPGLPGSPGHA-TEGPKGDRGPQGQPGLPGLPGPMGPPGLPGIDGVKGDKGNPGWPGAPGVPGPKGDPGFQGM 1291
α1(IV) 1182 KGDKGSKGEVGFP-GLAGSPGIPGSKGEQGFMGPPGPQGQPGLPGSPGHA-TEGPKGDRGPQGQPGLPGLPGPMGPPGLPGIDGVKGDKGNPGWPGAPGVPGPKGDPGFQGM 1291
α2(IV) 1225 PGERGDPGEANTLPGPVGVPGQKGDQGAPGERGPPGSPGLQGFPGITPPSNISGAPGDKGAPGIFGLKGYRGPPGPPGSAALPGSKGDTGNPGAPGTPGTKGWAGDSGPQGR 1336

α1(IV) 1292 PGIGGSPGITGSKGDMGPPGVPGFQGPKGLPGLQGIKGDQGDQGVPGAKGLPGPPGPPGPYDIIKGEPGLPGPEGPPGLKGLQGLPGPKGQQGVTGLVGIPGPPGIPGFDGA 1403
α1(IV) 1292 PGIGGSPGITGSKGDMGPPGVPGFQGPKGLPGLQGIKGDQGDQGVPGAKGLPGPPGPPGPYDIIKGEPGLPGPEGPPGLKGLQGLPGPKGQQGVTGLVGIPGPPGIPGFDGA 1403
α2(IV) 1337 PGVFGLPGEKGPRGEQGFMGNTGPTGAVGDRGPKGPKGDPGFPGAPGTVGAPGIAGIPQKIAVQPGTVGPQGRRGPPGAPGEMGPQGPPGEPGFRGAPGKAGPQGRGGVSAV 1448

α1(IV) 1404 PGQKGEMGPAGPTGPRGFPGPPGPDGLPGSMGPPGTPSVDHGFLVTRHSQTIDDPQCPSGTKILYHGYSLLYVQGNERAHGQDLGTAGSCLRKFSTMPFLFCNINNVCNFAS 1515
α1(IV) 1404 PGQKGEMGPAGPTGPRGFPGPPGPDGLPGSMGPPGTPSVDHGFLVTRHSQTIDDPQCPSGTKILYHGYSLLYVQGNERAHGQDLGTAGSCLRKFSTMPFLFCNINNVCNFAS 1515
α2(IV) 1449 PGFRGDEGPIGHQGPIGQEGAPGRPGSPGLPGMPGR-SVSIGYLLVKHSQTDQEPMCPVGMNKLWSGYSLLYFEGQEKAHNQDLGLAGSCLARFSTMPFLYCNPGDVCYYAS 1559

α1(IV) 1516 RNDYSYWLSTPEPMPMSMAPITGENIRPFISRCAVCEAPAMVMAVHSQTIQIPPCPSGWSSLWIGYSFVMHTSAGAEGSGQALASPGSCLEEFRSAPFIECHG-RGTCNYYA 1626
α1(IV) 1516 RNDYSYWLSTPEPMPMSMAPITGENIRPFISRCAVCEAPAMVMAVHSQTIQIPPCPSGWSSLWIGYSFVMHTSAGAEGSGQALASPGSCLEEFRSAPFIECHG-RGTCNYYA 1626
α2(IV) 1560 RNDKSYWLSTTAPLP--MMPVAEDEIKPYISRCSVCEAPAIAIAVHSQDVSIPHCPAGWRSLWIGYSFLMHTAAGDEGGGQSLVSPGSCLEDFRATPFIECNGGRGTCHYYA 1669

α1(IV) 1627 NAYSFWLATIERSEMFKKPTPSTLKAGELRTHVSRCQVCMRRT                                                                      1669
α1(IV) 1627 NAYSFWLATIERSEMFKKPTPSTLKAGELRTHVSRCQVCMRRT                                                                      1669
α2(IV) 1670 NKYSFWLTTIPEQSFQGSPSADTLKAGLIRTHISRCQVCMKNL                                                                      1712
```

### Col4 - $(\alpha5)_2\alpha6$

```
α5(IV)    1 --MKLRGVSLAAGLFLLALSLWGQPAEAAACYGCSPGSKCDCSGIKGEKGERGFPGLEGHPGLPGFPGPEGPPGPRGQKGDDGIPGPPGPKGIRGPPGLPGF  100
α5(IV)    1 --MKLRGVSLAAGLFLLALSLWGQPAEAAACYGCSPGSKCDCSGIKGEKGERGFPGLEGHPGLPGFPGPEGPPGPRGQKGDDGIPGPPGPKGIRGPPGLPGF  100
α6(IV)    1 MLINKLWLLLVTLCLTEELAAAGEKSYGKPCGGQDCSGSCQCFPEKGARGRPGPIGIQGPTGPQGFTGSTGLSGLKGERGFPGLLGPYGPKGDKGPMGVPGF  102

α5(IV)  101 PGTPGLPGMPGHDGAPGPQGIPGCNGTKGERGFPGSPGFPGLQGPPGPPGIPGMKGEPGSIIMSSLPGPKGNPGYPGPPGIQGLPGPTGIPGPIGPPGPPGL  202
α5(IV)  101 PGTPGLPGMPGHDGAPGPQGIPGCNGTKGERGFPGSPGFPGLQGPPGPPGIPGMKGEPGSIIMSSLPGPKGNPGYPGPPGIQGLPGPTGIPGPIGPPGPPGL  202
α6(IV)  103 LGINGIPGHPGQPGPRGPPGLDGCNGTQGAVGFPGPDGYPGLLGPPGLPGQKGSKGDP-VLAPGSFKGMKGDPGLPGLDGITGPQGAPGFPGAVGPAGPPGL  203

α5(IV)  203 MGPPGPPGLPGPKGNMGLNFQGPKGEKGEQGLQGPPGPPGQISEQKRPIDVEFQKGDQGLPGDRGPPGPPGIRGPPGPPG-GEKGEKGEQGEPGKRGKPGKD  303
α5(IV)  203 MGPPGPPGLPGPKGNMGLNFQGPKGEKGEQGLQGPPGPPGQISEQKRPIDVEFQKGDQGLPGDRGPPGPPGIRGPPGPPG-GEKGEKGEQGEPGKRGKPGKD  303
α6(IV)  204 QGPPGPPGPLGPDGNMGLGFQGEKGVKGDVGLPGPAGPPPSTGELEFMGFPKGKKGSKGEPGPKGFPGISGPPGFPGLGTTGEKGEKGEKGIPGLPGPRGPM  305

α5(IV)  304 GENGQPGIPGLPGDPGYPGEPGRDGEKGQKGDTGPPGPPGLV-IPRPGTGITIGEKGNIGLPGLPGEKGERGFPGIQGPPGLPGPPGAAVMGPPGPPGFPGE  404
α5(IV)  304 GENGQPGIPGLPGDPGYPGEPGRDGEKGQKGDTGPPGPPGLV-IPRPGTGITIGEKGNIGLPGLPGEKGERGFPGIQGPPGLPGPPGAAVMGPPGPPGFPGE  404
α6(IV)  306 GSEGVQGPPGQQGKKGTLGFPGLNGFQGIEGQKGDIGLPGPDVFIDIDGAVISGNPGDPGVPGLPGLKGDEGIQGLRGPSGVPGLP--ALSGVPGALGPQGF  405

α5(IV)  405 RGQKGDEGPPGISIPGPPGLDGQPGAPGLPGPPGPAGPHIPPSDEICEPGPPGPPGSPGDKGLQGEQGVKGDKGDT--CFNCIGTGISGPPGQPGLPGLPGP  504
α5(IV)  405 RGQKGDEGPPGISIPGPPGLDGQPGAPGLPGPPGPAGPHIPPSDEICEPGPPGPPGSPGDKGLQGEQGVKGDKGDT--CFNCIGTGISGPPGQPGLPGLPGP  504
α6(IV)  406 PGLKGDQGNPGRTTIGAAGLPGRDGLPGPPGPPGPPSPEFETETLHNKES--GFPGLRGEQGPKGNLGLKGIKGDSGFCACDGGVPNTGPPGEPGPPGPWGL  505

α5(IV)  505 PGSLGFPGQKGEKGQAGATGPKGLPGIPGAPGAPGFPGSKGEPGDILTFPGMKGDKGELGSPGAPGLPGLPGTPGQDGLPGLPGPKGEPGGITFKGERGPPG  606
α5(IV)  505 PGSLGFPGQKGEKGQAGATGPKGLPGIPGAPGAPGFPGSKGEPGDILTFPGMKGDKGELGSPGAPGLPGLPGTPGQDGLPGLPGPKGEPGGITFKGERGPPG  606
α6(IV)  506 IGLPGLKGARGDRGSGGAQGPAGAPGLVGPLGPSGPKGKKGEP-ILSTIQGMPGDRGDSGSQGFRGVIGEPGKDGVPGLPGLPGLPGD-GGQGFPGEKGLPG  605

α5(IV)  607 NPGLPGLPGNIGPMGPPGFGPPGPVGEKGIQGVAGNPGQPGIPGPKGDPGQTITQPGKPGLPGNPGRDGDVGLPGDPGLPGQPGLPGIPGSKGEPGIP-GIG  707
α5(IV)  607 NPGLPGLPGNIGPMGPPGFGPPGPVGEKGIQGVAGNPGQPGIPGPKGDPGQTITQPGKPGLPGNPGRDGDVGLPGDPGLPGQPGLPGIPGSKGEPGIP-GIG  707
α6(IV)  606 LPGEKGHPGPPGLPGNGLPGLPGPRGLPGDKGKDGLPGQQGLPGSKGIT-LPCIIPGSYGPSGFPGTPGFPGPKGSRGLPGTPGQPGSSGSKGEPGSPGLVH  706

α5(IV)  708 LPGPPGPKGFPGIPGPPGAPGTPGRIGLEGPPGPPGFPGPKGEPGFALPGPPGPPGLPGFKGALGPKGDRGFPGPPGPPGRTGLDGLPGPKGDVGPNGQPGP  809
α5(IV)  708 LPGPPGPKGFPGIPGPPGAPGTPGRIGLEGPPGPPGFPGPKGEPGFALPGPPGPPGLPGFKGALGPKGDRGFPGPPGPPGRTGLDGLPGPKGDVGPNGQPGP  809
α6(IV)  707 LPELPGFPGPRGEKGLPGFPGLPGKDGLPGMIGSPGLPGSKGATGDIFGAENGAPGEQGLQGLTGHKGFLGDSGLPGLKGVHGKPGLLGPKGERGSPGTPGQ  808

α5(IV)  810 MGPPGLP-GIGVQGPPGPPGIPGPIGQPGLHGIPGEKGDPGPPGLD----VPGPPGERGSPGIPGAPGPIGPPGSPGLPGKAGASGFPGTKGEMGMMGPPGP  906
α5(IV)  810 MGPPGLP-GIGVQGPPGPPGIPGPIGQPGLHGIPGEKGDPGPPGLD----VPGPPGERGSPGIPGAPGPIGPPGSPGLPGKAGASGFPGTKGEMGMMGPPGP  906
α6(IV)  809 VGQPGTPGSSGPYGIKGKSGLPGAPGFPGISGHPGKKGTRGKKGPPGSIVKKGLPGLKGLPGNPGLVGLKGSPGSPGVAGLPALSGPKGEKGSVGFVGFPGI  910

α5(IV)  907 PGPLGIPGRSGVPGLKGDDGLQGQPGLPGPTGEKGSKGEPGLPGPPGPM-DPNLLGSKGEKGEPGLPGIPGVSGPKGYQGLPGDPGQPGLSGQPGLPGPP-G 1006
α5(IV)  907 PGPLGIPGRSGVPGLKGDDGLQGQPGLPGPTGEKGSKGEPGLPGPPGPM-DPNLLGSKGEKGEPGLPGIPGVSGPKGYQGLPGDPGQPGLSGQPGLPGPP-G 1006
α6(IV)  911 PGLPGIPGTRGLKGIPGSTGKMGPSGRAGTPGEKGDRGNPGPVGIPSPRRPMSNLWLKGDKGSQGSAGSNGFPGPRGDKGEAGRPGPPGLPGAPGLPGIIKG 1012

α5(IV) 1007 PKGNPGLPGQPGLIGPPGLKGTIGDMGFPGPQGVEGPPGPSGVPGQPGSPGLPGQKGDKGDPGISSIGLPGLPGPKGEPGLPGYPGNPGIKGSVGDPGLPGL 1108
α5(IV) 1007 PKGNPGLPGQPGLIGPPGLKGTIGDMGFPGPQGVEGPPGPSGVPGQPGSPGLPGQKGDKGDPGISSIGLPGLPGPKGEPGLPGYPGNPGIKGSVGDPGLPGL 1108
α6(IV) 1013 VSGKPGPPGFMGIRGLPGLKGSSGITGFPGMPGESGSQGIRGSPGLPGASGLPGLKGDNGQT----VEISGSPGPKGQPGESGFKGTKGRDGLIGNIGFPGN 1110

α5(IV) 1109 PGTPGAKGQPGLPGFPGTPGPPGPKGISGPPGNPGLPGEPGPVGGGGHPGQPGPPGEKGKPGQDGIPGPAGQKGEPGQPGFGNPGPPGLPGLSGQKGDGGLP 1210
α5(IV) 1109 PGTPGAKGQPGLPGFPGTPGPPGPKGISGPPGNPGLPGEPGPVGGGGHPGQPGPPGEKGKPGQDGIPGPAGQKGEPGQPGFGNPGPPGLPGLSGQKGDGGLP 1210
α6(IV) 1111 KGEDGKVGVSGDVGLPGAPGFPGVAGMRGEPGLPGSSGHQGAIGPLGSPGLIGPKGFPGFPGLHGLNGLPGTKGTHGTPGPSITGVPGPAGLPGPKGEKGYP 1212

α5(IV) 1211 GIP-GNPGLPGPKGEPGFHGFPGVQGPPGPPGSPGPA----LEGPKGNPGPQGPPGRPGLPGPEGPPGLPGNGGIKGEKGNPGQPGLPGLPGLKGDQGPPGL 1307
α5(IV) 1211 GIP-GNPGLPGPKGEPGFHGFPGVQGPPGPPGSPGPA----LEGPKGNPGPQGPPGRPGLPGPEGPPGLPGNGGIKGEKGNPGQPGLPGLPGLKGDQGPPGL 1307
α6(IV) 1213 GIGIGAPGKPGLRGQKGDRGFPGLQGPAGLPGAPGISLPSLIAGQPGDPGRPGLDGERGRPGPAGPPGPPGPSSNQGDTGDPGFPGIPGPKGPKGDQGIPGF 1314

α5(IV) 1308 QGNPGRPGLNGMKGDPGLPGVPGFPGMKGPSGVPGSAGPEGEPGLIGPPGPPGLPGPSGQSIIIKGDAGPPGIPGQPGLKGLPGPQGPQGLPGPTGPPGDPG 1409
α5(IV) 1308 QGNPGRPGLNGMKGDPGLPGVPGFPGMKGPSGVPGSAGPEGEPGLIGPPGPPGLPGPSGQSIIIKGDAGPPGIPGQPGLKGLPGPQGPQGLPGPTGPPGDPG 1409
α6(IV) 1315 SGLPGELGLKGMRGEPGFMGTPGKVGPPGDPGFPGMKGKAGPRGSSGLQGDPGQTPTAEAVQVPPGPLGLPGIDGIPGLTGDPGAQGPVGLQGSKGLPGIPG 1416

α5(IV) 1410 RNGLPGFDGAGGRKGDPGLPGQPGTRGLDGPPGPDGLQGPPGPPGTSSVAHGFLITRHSQTTDAPQCPQGTLQVYEGFSLLYVQGNKRAHGQDLGTAGSCLR 1511
α5(IV) 1410 RNGLPGFDGAGGRKGDPGLPGQPGTRGLDGPPGPDGLQGPPGPPGTSSVAHGFLITRHSQTTDAPQCPQGTLQVYEGFSLLYVQGNKRAHGQDLGTAGSCLR 1511
α6(IV) 1417 KDGPSGLPGPPGALGDPGLPGLQGPPGFEGAPGQQGPFGMPGMPGQS-MRVGYTLVKHSQSEQVPPCPIGMSQLWVGYSLLFVEGQEKAHNQDLGFAGSCLP 1517

α5(IV) 1512 RFSTMPFMFCNINNVCNFASRNDYSYWLSTPEPMPMSMQPLKGQSIQPFISRCAVCEAPAVVIAVHSQTIQIPHCPQGWDSLWIGYSFMMHTSAGAEGSGQA 1613
α5(IV) 1512 RFSTMPFMFCNINNVCNFASRNDYSYWLSTPEPMPMSMQPLKGQSIQPFISRCAVCEAPAVVIAVHSQTIQIPHCPQGWDSLWIGYSFMMHTSAGAEGSGQA 1613
α6(IV) 1518 RFSTMPFIYCNINEVCHYARRNDKSYWLSTTAPIPM-MPVSQTQIPQYISRCSVCEAPSQAIAVHSQDITIPQCPLGWRSLWIGYSFLMHTAAGAEGGGQSL 1618

α5(IV) 1614 LASPGSCLEEFRSAPFIECH-GRGTCNYYANSYSFWLATVDVSDMFSK-PQSETLKAGDLRTRISRCQVCMKRT                             1685
α5(IV) 1614 LASPGSCLEEFRSAPFIECH-GRGTCNYYANSYSFWLATVDVSDMFSK-PQSETLKAGDLRTRISRCQVCMKRT                             1685
α6(IV) 1619 VSPGSCLEDFRATPFIECSGARGTCHYFANKYSFWLTTVEERQQFGELPVSETLKAGQLHTRVSRCQVCMKSL-                             1691
```

### Col4 - ($\alpha3\alpha4\alpha5$)

```
α3(IV)    1 ----------------MSARTAPRPQVLLLPLLLVLLAAAPAASKGCVCKDKGQCFCDGAKGEKGEKGFPGPPGSPGQKGFTGPEGLPGPQGPKGFPGLPGLTGSKGVRGISGLP   99
α4(IV)    1 MWSLHIVLMRCSFRLTKSLATGPWSLILILFSVQYVYGSGKKYIGPCGGRDCSVCHCVPEKGSRGPPGPPGPQGPIGPLGAPGPIGLSGEKGMRGDRGPPGAAGDKGDKGPTGVP  115
α5(IV)    1 -----------------MKLRGVSLAAGLFLLALSLWGQPAEAAACYGCSPGSKCDCSGIKGEKGERGFPGLEGHPGLPGFPGPEGPPGPRGQKGDDGIPGPPGPKGIRGPPGLP   98

α3(IV)  100 GFSGSPGLPGTPGNTGPYGLVGVPGCSGSKGEQGFPGLPGTLGYPGIPGAAGLKGQKGAP-AKEEDIELDAKGDPGLPGAPGPQGLPGPPGFPGPVGPPGPPGFFGFPGAMGPRG  213
α4(IV)  116 GFPGLDGIPGHPGPPGPRGKPGMSGHNGSRGDPGFPGGRGALGPGGPLGHPGEKGEKGNS-VFILGAVKGIQGDRGDPGLPGLPGSWGAGGPAGPTGYPGEPGLVGPPGQPGRPG  229
α5(IV)   99 GFPGTPGLPGMPGHDGAPGPQGIPGCNGTKGERGFPGSPGFPGLQGPPGPPGIPGMKGEPGSIIMSSLPGPKGNPGYPGPPGIQGLPGPTGIPGPIGPPGPPGLMGPPGPPGLPG  213

α3(IV)  214 PKGHMGERVIGHKGERGVKGLTGPPGPPGTVIVTLTGPDNRTDLKGEKGDKGAMGEPGPPGPSGLPGES--YGSEKGAPGDPGLQGKPGKDGVPGFPGSEGVKGNRGFPGLMGED  326
α4(IV)  230 LKGNPGVGVKGQMGDPGEVGQQGSPGPT--LLVEPPDFCLYKGEKGIKGIPGMVGLPGPPGRKGESGIG--AKGEKGIPGFPGPRGDPGSYGSPGFPGLKGELGLVGDPGLFGLI  340
α5(IV)  214 PKGNMGLNFQGPKGEKGEQGLQGPPGPPGQI-SEQKRPIDVEFQKGDQGLPGDRGPPGPPGIRGPPGPPGGEKGEKGEQGEPGKRGKPGKDGENGQPGIPGLPGDPGYPGEPGRD  327

α3(IV)  327 GIKGQKGDIGPPGFRGPTEYYD----TYQEKGDEGTPGPPGPRGARGPQGPSGPPGVPGSPGSS---RPGLRGAPGWPGLKGSKGERGRPGKD--AMGTPGSPGCAGSPGLPGSP  432
α4(IV)  341 GPKGDPGNRGHPGPPGVLVTPP-----LPLKGPPGDPGFPGRYGETGDVGPPGPPGLLGRPGEACAGMIGPPGPQGFPGLPGLPGEAGIPGRPDSAPGKPGKPGSPGLPGAPGLQ  450
α5(IV)  328 GEKGQKGDTGPPGPPGLVIPRPGTGITIGEKGNIGLPGLPGEKGERGFPGIQGPPGLPGPPGAAVM---GPPGPPGFPGERGQKGDEGPPGIS--IPGPPGLDGQPGAPGLPGPP  437

α3(IV)  433 GPPGPPGDIVFRKGPPGDHGLPGYLGSPGIPGVDGPKGEPGLLCTQCPYIP--GPPGLPGLPGLHGVKGIPGRQGAAGLKGSPGSPGNTGLPGFPGFPGAQGDPGLKGEKGETLQ  545
α4(IV)  451 GLPGSS-VIYCSVGNPGPQGIKGKVGPPGGRGPKGEKGNEGL-C-ACEP-GPMGPPGPPGLPGRQGSKGDLGLPGWLGTKGDPGPPGAEGPPGLPGKHGASGPPGNKGAKGDM-V  560
α5(IV)  438 GPAGPHIPPSDEICEPGPPGPPGSPGDKGLQGEQGVKGDKGDTCFNCIGTGISGPPGQPGLPGLPGPPGSLGFPGQKGEKGQAGATGPKGLPGIPGAPGAPGFPGSKGEPGDILT  552

α3(IV)  546 PEGQVGVPGDPGLRGQPGRKGLDGIPGTPGVKGLPGPKGELALSGEKGDQGPPGDPGSPGSPGPAGPAGPPGYGPQGEPGLQGTQGVPGAPGPPGEAGPRGEL----SVSTPVPG  656
α4(IV)  561 VSRVKGHKGERGPDGPPGFPGQPGSHGRDGHAGEKGDPGPPGDH-EDATPGGKGFPGPLGPPGKAGPVGPPGLGFPGPPGERGHPGVPGHPGVRGPDGLKGQKGDTISCNVTYPG  674
α5(IV)  553 FPGMKGDKGELGSPGAPGLPGLPGTPGQDGLPGLPGPKGEPGGITFKGERGPPGNPGLPGLPGNIGPMGPPGFGPPGPVGEKGIQGVAGNPGQPGIPGPKGDPGQT----ITQPG  663

α3(IV)  657 PPGPPGPPGHPGPQGPPGIPGSLGKCGDPGLPGPDGEPGIPGIGFPGPPGPKGDQGFPGTKGSLGCPGKMGEPGLPGKPGLPGAKGEP---AVAMPGGPGTPGFPGERGNSGEHG  768
α4(IV)  675 RHGPPGFDGPPGPKGFPGPQGAPGLSGSDGHKGRPGTPGTA-E-IPGPPGFRGDMGDPGFGGEKGSS-PVGPPGPPGSPGVNGQKGIPGDPAFGHLGPPGKRGLSGVPGIKGPRG  786
α5(IV)  664 KPGLPGNPGRDGDVGLPGDPGLPGQPGLPGIPGSKGEPGIPGIGLPGPPGPKGFPGIPGPPGAPGTPGRIGLEGPPGPPGFPGPKGEPGFA---LPGPPGPPGLPGFKGALGPKG  775

α3(IV)  769 EIGLPGLPGLPGTPGNEGLDGPRGDPGQPGPPGEQGPPGRC-IEGPRGAQGLPGLNGLKGQQGRRGKTGPKGDPGIPGLDRSGFPGETGSPGIPGHQGEMGPLGQRGYPGNPGIL  882
α4(IV)  787 DPGCPGAEGPAGIPGFLGLKGPKGREGHAGFPGVPGPPGHSCERGAPGIPGQPGLPGYPGSPGAPGGKGQPGDVGPPGPA--GMKGLPGLPGRPGAHGPPGLPGIPGPFGDDGLP  899
α5(IV)  776 DRGFPGPPGPPGRTGLDGLPGPKGDVGPNGQPGPMGPPGLP-GIGVQGPPGPPGIPGPIGQPGLHGIPGEKGDPGPPGLDVPGPPGERGSPGIPGAPGPIGPPGSPGLPGKAGAS  889

α3(IV)  883 GPPGEDGVIGMMGFPGAIGPPGPPGNPGTPGQRGSPGIPGVKGQRGTPGAKGEQGDKGNPGPS-----EISHVIGDKGEPGLKGFAGNPGEKGNRGVPGMPGLKGLKGLPGPAGP  992
α4(IV)  900 GPPGPKGPRGLPGFPGFPGERGKPGAEGCPGAKGEPGEKGMSGLPGDRGLRGAKGAIGPPGDEGEM-AIISQKGTPGEPGPPGDDGFPGERGDKGTPGMQGRRGEPGRYGPPGFH 1013
α5(IV)  890 GFPGTKGEMGMMGPPGPPGPLGIPGRSGVPGLKGDDGLQGQPGLPGPTGEKGSKGEPGLPGPPGPMDPNLLGSKGEKGEPGLPGIPGVSGPKGYQGLPGDPGQPGLSGQPGLPGP 1004

α3(IV)  993 P-GPRGDLGSTGNPGEPGLRGIPGSMGNMGMPGSKGKRGTLGFPGRAGRPGLPGIHGLQGDKGEPGYSEGTRPGPPGPTGDPGLPGDMGKKGEMGQPGPPGHLGPAGPEGAPGSP 1106
α4(IV) 1014 RGEPGEKGQPGPPGPPGPPGSTGLRGFIGFPGLPGDQGE-PGSPGPPGFSGIDGARGPKGNKGDP-ASHFGPPGPKGEPGSPGCPGHFGASGEQGLPGIQGPRGSPGRPGPPGSS 1126
α5(IV) 1005 P-GPKGNPGLPGQPGLIGPPGLKGTIGDMGFPGPQGVEGPPGPSGVPGQPGSPGLPGQKGDKGDPGISSIGLPGLPGPKGEPGLPGYPGNPGIKGSVGDPGLPGLPGTPGAKGQP 1118

α3(IV) 1107 GSPGLPGKPGPHGDLGFKGIKGLLGPPGIRGPPGLPGFPGSPGPMGIRGDQGRDGIPGPAGEKGETGLL-RAPPGPRGNPGAQGAKGDRGAPGFP--GLPGRKGAMGDAGPRGPT 1218
α4(IV) 1127 GPPGCPGDHGMPGLRGQPGEMGDPGPRGLQGDPGIPGPPGIKGPSGSPGLNGLHGLKGQKGTKGASGLHDVGPPGPVGIPGLKGERGDPGSPGISPPGPRGKKGPPGPPGSSGPP 1241
α5(IV) 1119 GLPGFPGTPGPPGPKGISGPPGNPGLPGEPGPVGGGGHPGQPGPPGEKGKPGQDGIPGPAGQKGEPGQPGFGNPGPPGLPGLSGQKGDGGLPGIP--GNPGLPGPKGEPGFHGFP 1231

α3(IV) 1219 GIEGFPGPPGLPGAIIPGQTGNRGPPGSRGSPGAPGPPGPPGSH-VIGIKGDKGSMGHPGPKGPPGTAGDMGPPGRLGAPGTPGLPGPRGDPGFQGFPGVKGEKGNPGFLGSIGP 1332
α4(IV) 1242 GPAGATGRA-PKDIPDPGPPGDQGPPGPDGPRGAPGPPGLPGSV-DLLR-GEPGDCGLPGPPGPPGPPGPPGYKGFPGCDGKDGQKGPVGFPGPQGPHGFPGPPGEKGLPGPPGR 1353
α5(IV) 1232 GVQGPPGPPGSPGPALEGPKGNPGPQGPPGRPGLPGPEGPPGLPGNGGIKGEKGNPGQPGLPGLPGLKGDQGPPGLQGNPGRPGLNGMKGDPGLPGVPGFPGMKGPSGVPGSAGP 1346

α3(IV) 1333 PGPIGPKGPPGVRGDPGTL-KIISLPGSPGPPGTPGEPGMQGEPGPPGPPGNLGPCGPRGKPGKDGKPGTPGPAGEKGNKGSKGEPGPAGSDGLPGLKGKRGDSGSP-ATWTTRG 1445
α4(IV) 1354 KGPTGLPGPRGEPGPPADVDDCPRIPGLPGAPGMRGPEGAMGLPGMRGPSGP-GCKGEPGLDGRRGVDGVPGSPGPPGRKGDTGEDGYPGGPGPPGPIGDPGPKGF-G-PGYLGG 1465
α5(IV) 1347 EGEPGLIGPPGPPGLPGPSGQSIIIKGDAGPPGIPGQPGLKGLPGPQGPQGLPGPTGPPGDPGRNGLPGFDGAGGRKGDPGLPGQPGTRGLDGPPGPDGLQGPPGPPGTSSVAHG 1461

α3(IV) 1446 FVFTRHSQTTAIPSCPEGTVPLYSGFSFLFVQGNQRAHGQDLGTLGSCLQRFT--PFLFCNVNDVCNFASRNDYSYWLSTPALMPMNMAPITGRALEPYISRCTVCEGPAIAIAV 1558
α4(IV) 1466 FLLVLHSQTDQEPTCPLGMPRLWTGYSLLYLEGQEKAHNQDLGLAGSCLPVFSTLPFAYCNIHQVCHYAQRNDRSYWLASAAPLPM--MPLSEEAIRPYVSRCAVCEAPAQAVAV 1578
α5(IV) 1462 FLITRHSQTTDAPQCPQGTLQVYEGFSLLYVQGNKRAHGQDLGTAGSCLRRFSTMPFMFCNINNVCNFASRNDYSYWLSTPEPMPMSMQPLKGQSIQPFISRCAVCEAPAVVIAV 1576

α3(IV) 1559 HSQTTDIPPCPHGWISLWKGFSFIMFTSAGSEGTGQALASPGSCLEEFRASPFLECHG-RGTCNYYSNSYSFWLASLNPERMFRK-PIPSTVKAGELE-KIISRCQVCMKKRH   1668
α4(IV) 1579 HSQDQSIPPCPQTWRSLWIGYSFLMHTGAGDQGGGQALMSPGSCLEDFRAAPFLECQGRQGTCHFFANKYSFWLTTVKADLQFSSAPAPDTLKESQAQRQKISRCQVCVKYS-   1690
α5(IV) 1577 HSQTIQIPHCPQGWDSLWIGYSFMMHTSAGAEGSGQALASPGSCLEEFRSAPFIECHG-RGTCNYYANSYSFWLATVDVSDMFSK-PQSETLKAGDLR-TRISRCQVCMKRT-   1685
```

**Supplementary Fig. 2.** Alignment of Col10 $(\alpha1)_3$ homotrimer and the Col4 heterotrimers $(\alpha1)_2\alpha2$, $(\alpha5)_2\alpha6$ and $\alpha3\alpha4\alpha5$ indicating the triple-helical domain (grey) and sites that we believe could represent potential DDR-binding sites (purple/blue/orange). It should be noted that these alignments simply show the relative position of the $\alpha$-chains in the triple helix and do not represent a multiple sequence alignment (MSA). Further, the perfect Gly-X-Y repeats in the Col4 $\alpha$-chains are frequently interspersed with non-helical interruptions whose lengths might also vary between the three chains. Structural consequence of these interruptions and how they might influence the start and ends of the triple-helical domains on either side is currently not clear. To circumvent this, the interruptions have previously been padded with gaps, represented by dashes "-", such that the triple-helical length is optimal on either side.[2] Thus, dashes here carry a different meaning than those in conventional MSAs.

**Supplementary Table 1.** Peptides used in this study contain three primary sequence features – KGE motifs on the C- and N-terminal to facilitate covalent capture (orange), DDR recognition motif (cyan), C-terminal Tyr for accurate spectrophotometric determination of concentration (green) and Lys (blue) or Asp (red) to guide the binding specificity. Theoretical molecular weight of the peptide monomers was calculated from their amino acid sequences. The expected molecular weight of the corresponding triple-helical trimers was calculated assuming assembly of three peptide monomers and were compared with the experimentally observed molecular masses obtained by MALDI–TOF mass spectrometry.

| acronym | sequence | Molecular weight, monomer (Da) | Molecular weight, trimer (Da) |
|---|---|---|---|
| pep-P-cc | Ac–GPP–GP**KGE**P–GPP–GP**R**–GPP–GV(Nle)GFO–(GPP)$_3$–GP**KGE**P–GPP–GG**Y** | 4123.6 | 12370.8 |
| pep-K-cc | Ac–GPP–GP**KGE**P–GPP–GP**R**–GPP–GV(Nle)GFO–GP**K**(GPP)$_2$–GP**KGE**P–GPP–GG**Y** | 4154.6 | 12463.8 |
| pep-D-cc | Ac–GPP–GP**KGE**P–GPP–GP**R**–GPP–GV(Nle)GFO–GP**D**(GPP)$_2$–GP**KGE**P–GPP–GG**Y** | 4141.6 | 12424.8 |

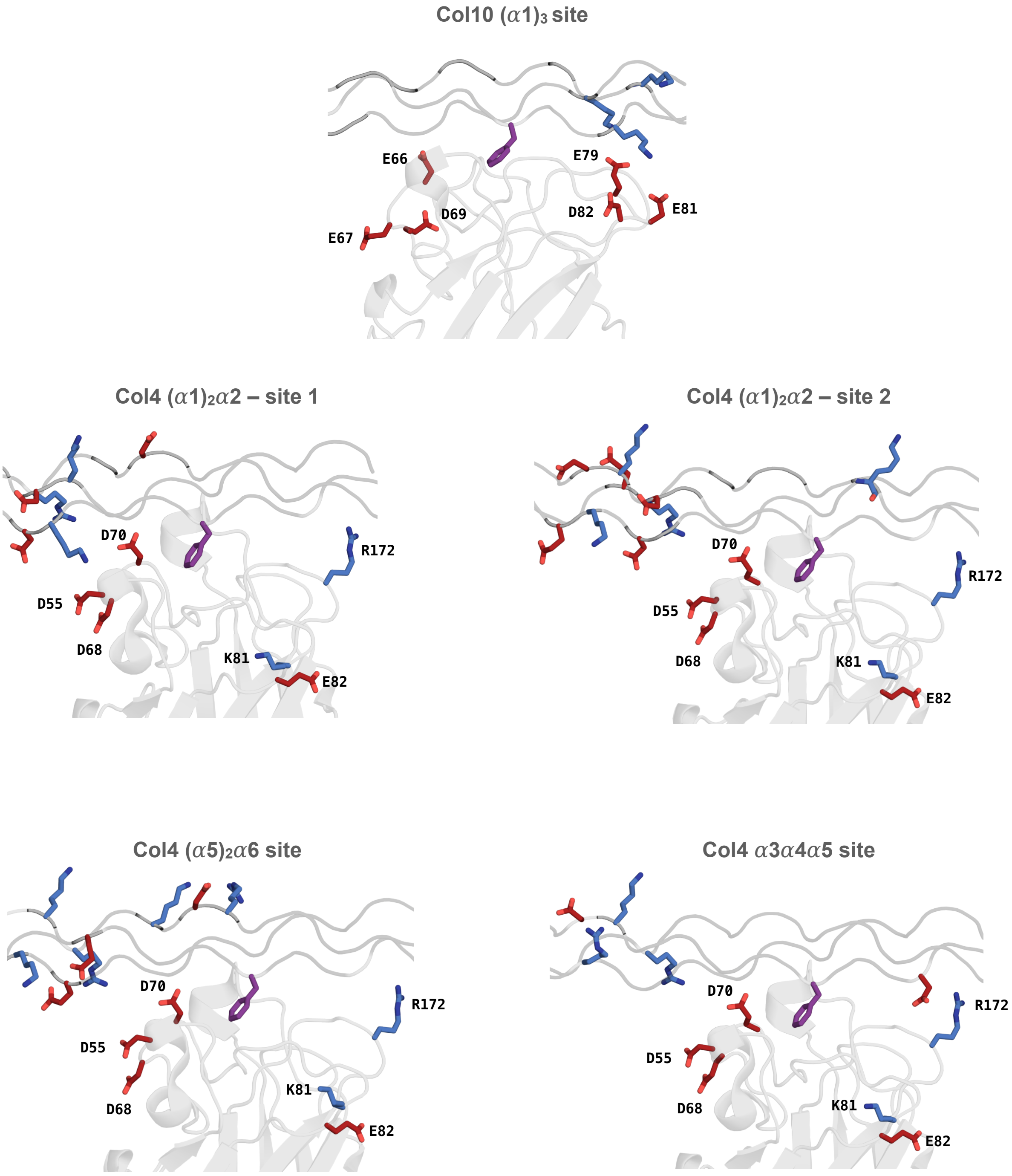


**Supplementary Fig. 3.** Proposed interactions between the positive (red) and negatively (blue) charged residues of the potential binding sites identified in Col10 and Col4 heterotrimers with the DS-domains of DDR2 and DDR1, respectively. The images were created by mutating the sequence positions in PyMOL to those observed in the potential binding sites, using the previously published crystal structure of DDR2 (pdb: 2wuh) as a template. Phe of the middle chain bound to the DS-domains is shown in purple.

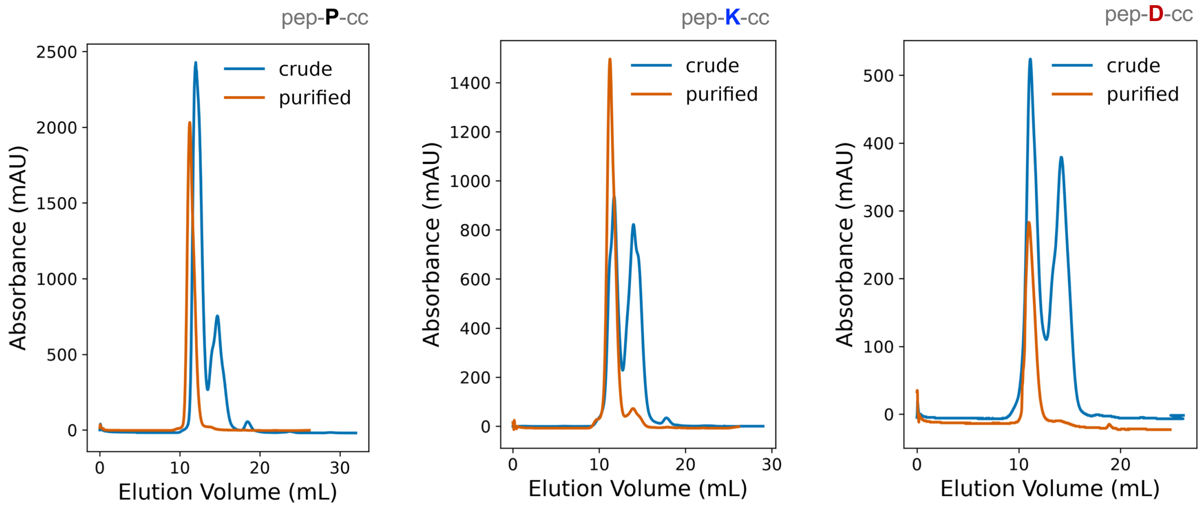


**Supplementary Fig. 4.** Size-exclusion chromatograms (SEC) of crude (blue) and purified (orange) covalently captured constructs pep-P-cc, pep-K-cc and pep-D-cc recorded at 280 nm. Fractions eluting from the SEC run of the crude reaction mixture were analyzed using MALDI-TOF mass spectrometry and those containing the desired construct were pooled and run again to obtain the chromatograms shown as purified constructs. The fractions obtained from the SEC run of the purified construct were used for all subsequent biophysical characterization and DDR binding assays. This two-step procedure was followed to ensure high purity. The chromatograms are representative of two independent reactions.

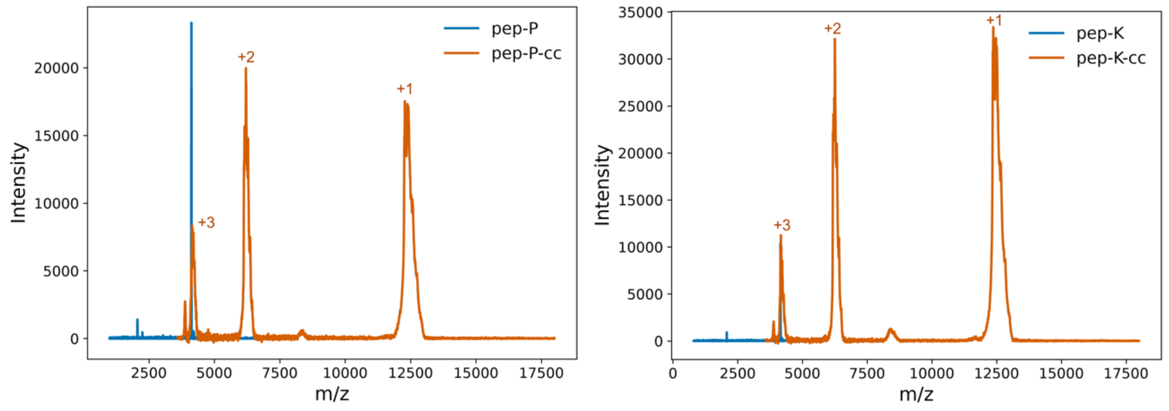

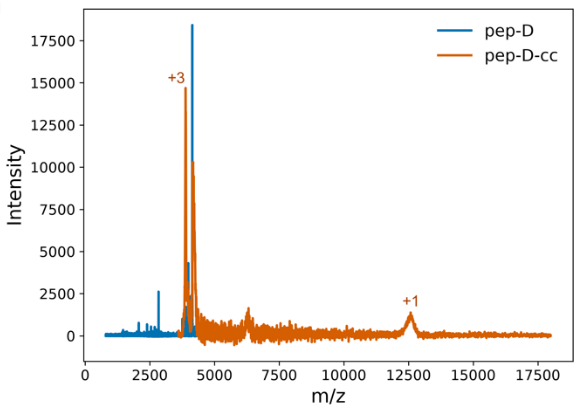


**Supplementary Fig. 5.** MALDI–TOF mass spectra of native peptide pep-P, pep-K and pep-D and their covalently captured counterparts pep-P-cc, pep-K-cc and pep-D-cc. The native peptides show a single peak corresponding to the expected mass while the covalently captured products exhibit multiply charged peaks in addition.

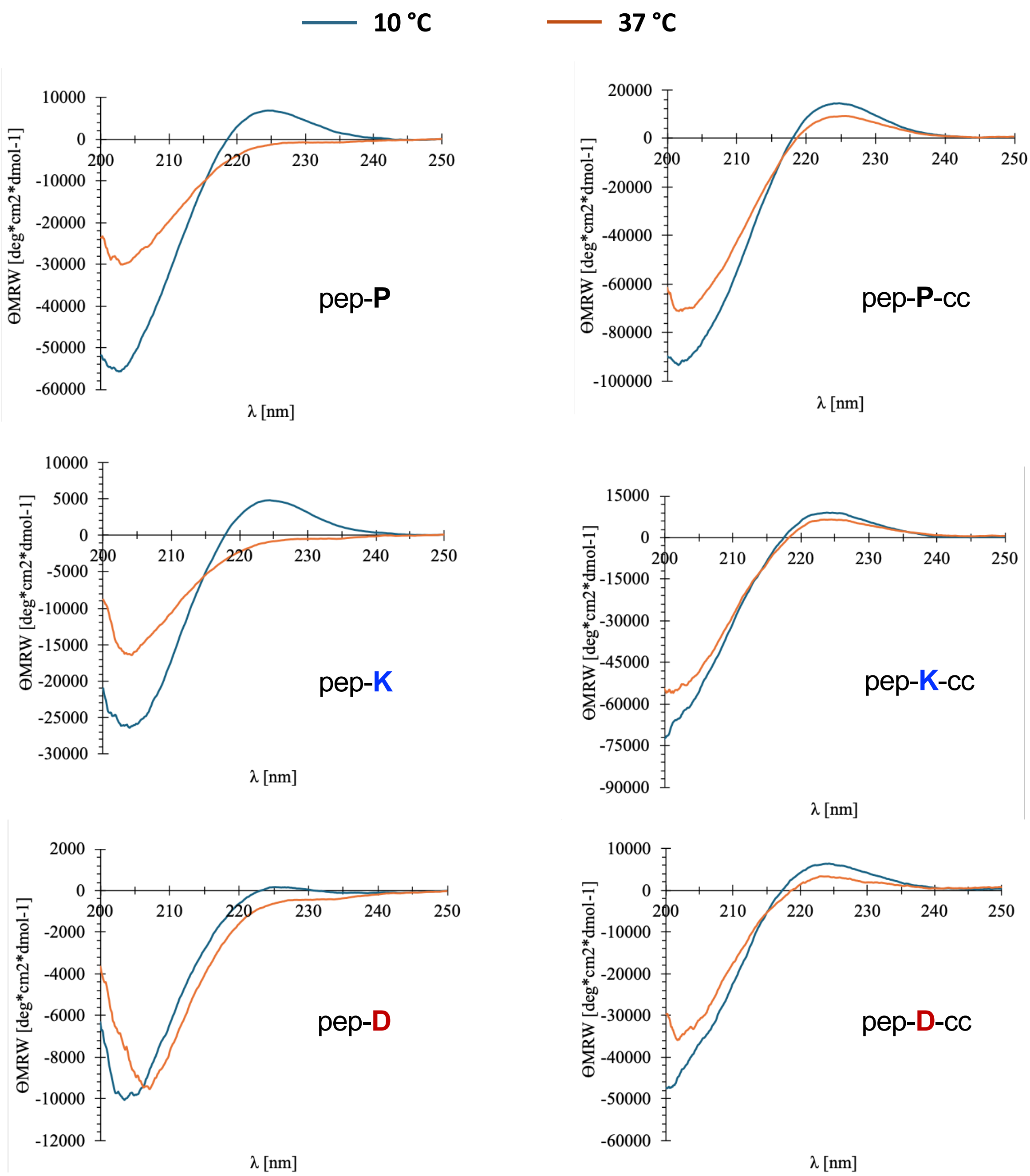


**Supplementary Fig. 6.** Circular dichroism (CD) spectra of native (blue) and covalently captured (orange) triple-helical peptides. CD spectra of pep-P, pep-K, pep-D (left panel) and their covalently captured constructs pep-P-cc, pep-K-cc and pep-D-cc (right panel) were recorded at 10 °C and after unfolding at 37 °C. The raw ellipticity obtained was converted to molar residue weight (MRW) shown on the Y-axis. The native triple helices unfold completely at 37°C as suggested by the loss in positive signal at 225 nm with the exception of pep-D, which appears to remain unfolded even at 10°C. Thus, pep-D appears to have lower stability than pep-P or pep-K. In contrast, the covalently captured constructs retain the positive maximum for all peptides suggesting that the covalent linkage increase the stability of the triple helix and prevents it from unfolding.

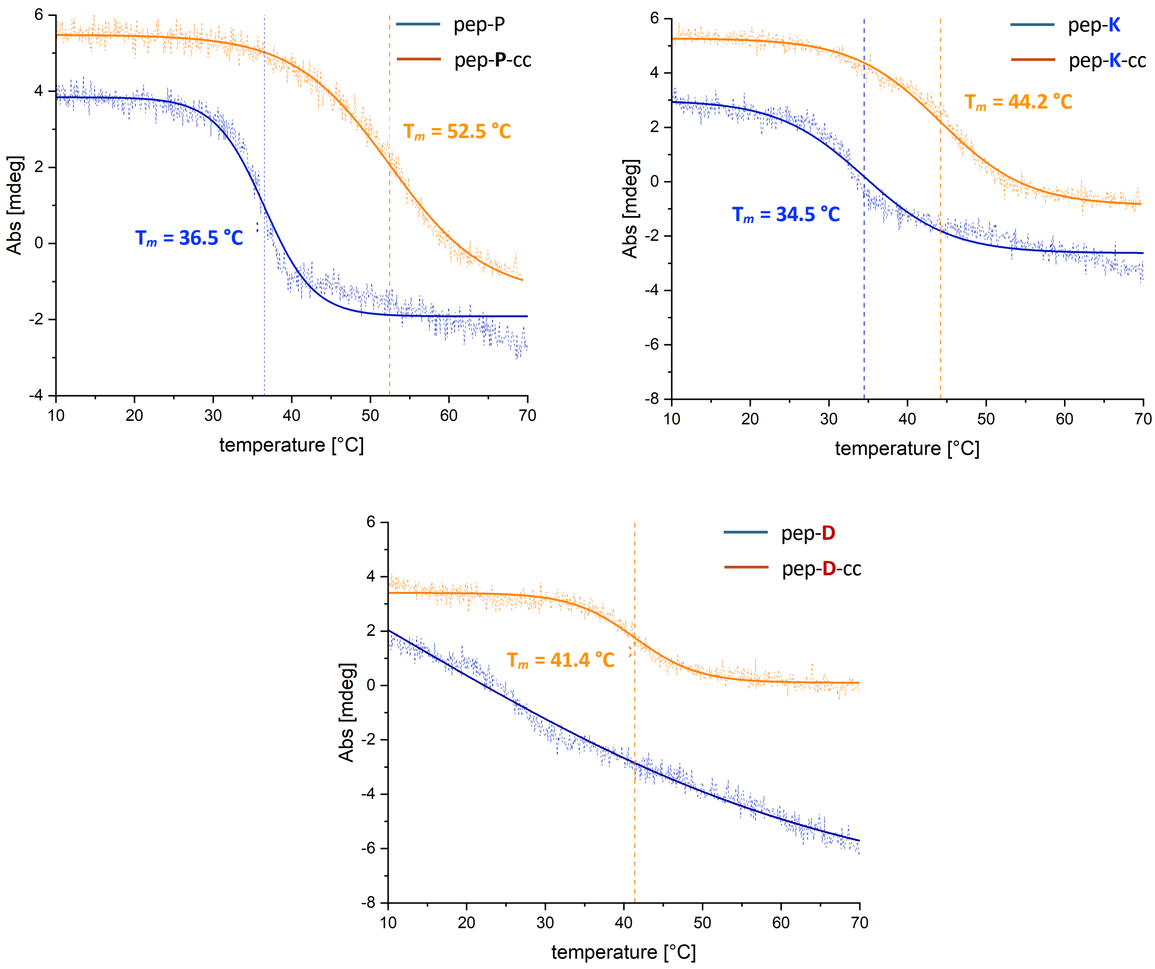


**Supplementary Fig. 7.** Thermal unfolding of native and covalently captured peptides monitored by CD spectroscopy. Thermal denaturation curves before and after covalent capture were recorded by monitoring the CD signal at 225 nm over the indicated temperature range. Solid lines represent nonlinear fits to the experimental data. The melting temperatures ($T_m$), defined as the inflection point of the fitted transition, are indicated by the dashed vertical lines.

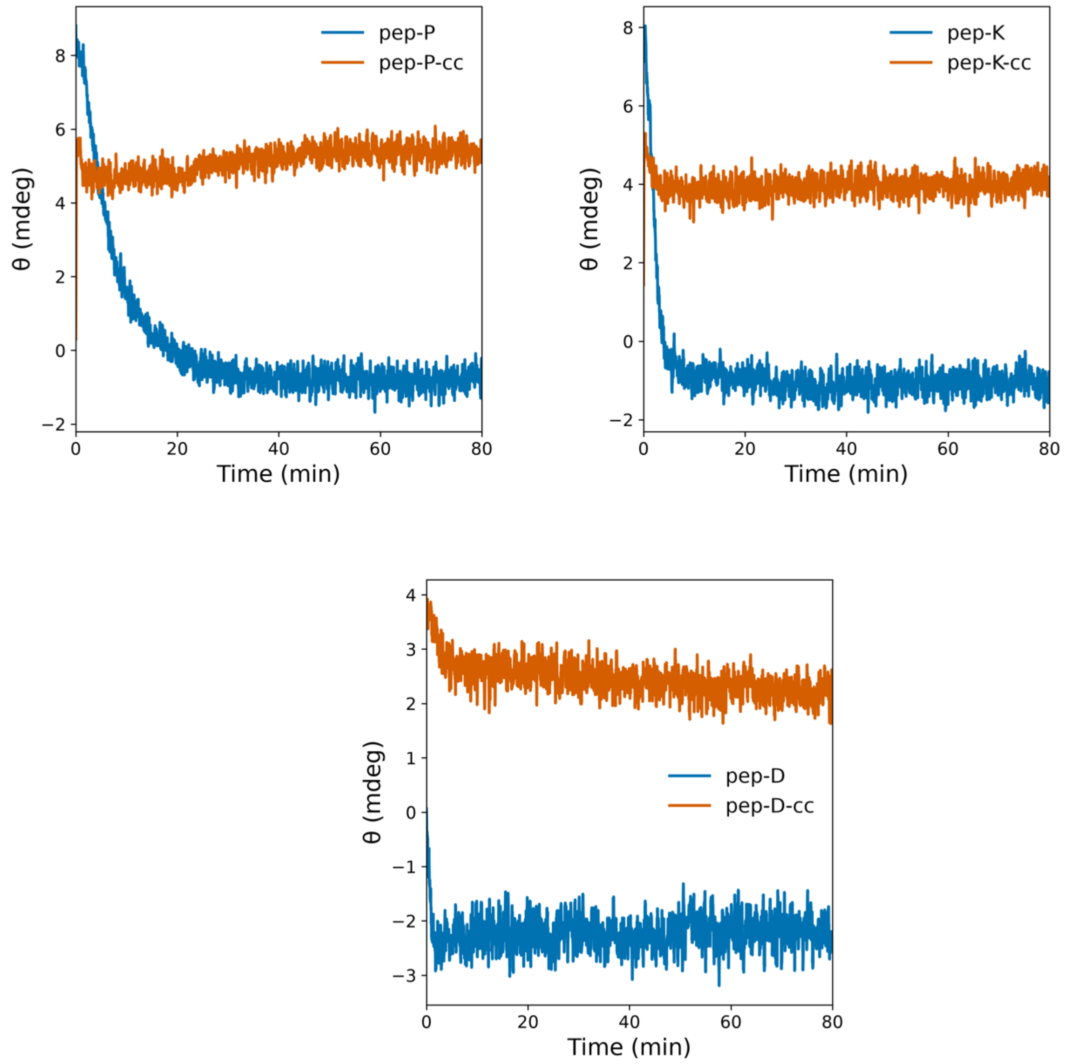


**Supplementary Fig. 8.** Comparison of the time-dependent change in ellipticity at 225 nm as recorded in CD spectroscopy for native (pep-P, pep-K and pep-D) and covalently captured peptides (pep-P-cc, pep-K-cc and pep-D-cc) at 37°C. The native peptides exhibited a rapid loss of the characteristic collagen triple-helical signal, whereas the covalently captured peptides maintained stable ellipticity throughout the duration of the measurement, demonstrating the ability to retain the folded triple-helical structure at physiological temperature. In the case of pep-D, the signal shows a dramatic loss upon transferring to 37°C indicating comparatively lower stability than pep-P or pep-K, both of which show a gradual decrease in ellipticity.
